# GCNA is a histone binding protein required for spermatogonial stem cell maintenance

**DOI:** 10.1101/2022.02.21.481287

**Authors:** Jonathan Ribeiro, Gerry P. Crossan

## Abstract

Recycling and *de-novo* deposition of histones during DNA replication is a critical challenge faced by eukaryotic cells and is coordinated by histone chaperones. However, little is known about how tissue-specific histone chaperones function to maintain tissue homeostasis. Here we show that Germ Cell Nuclear Acidic protein (GCNA), a germ cell specific protein in adult mice, can bind histones and purified GCNA exhibits histone chaperone activity. GCNA associates with the DNA replication machinery and supports progression through S-phase in murine spermatogonial stem cells (SSCs). Whilst GCNA is dispensable for embryonic germ cell development, it is required for the maintenance of the SSC pool and for long-term production of sperm. Our work describes the role of a germ cell specific histone chaperone in SSCs maintenance in mice. These findings provide a mechanistic basis for the male infertility observed in patients carrying mutations in the *GCNA* locus.

## Introduction

The germline is tasked with faithfully transmitting genetic information from one generation to the next. The earliest mammalian germ cells, Primordial Germ Cells (PGCs), emerge during early embryonic development and migrate to the genital ridges (Tang et al., 2016). In the embryonic gonad, these cells undergo rapid proliferation and extensive epigenetic changes. Subsequently, female embryonic germ cells initiate meiosis, a process that does not begin until postnatal life in males. The male embryonic germ cells, in contrast, give rise to a population of self-renewing spermatogonial stem cells (SSCs)(Griswold, 2021). At the onset of puberty, SSCs resume proliferation, differentiate and initiate spermatogenesis. The capacity of SSCs to self-renew underlies the ability of males to produce gametes throughout their entire lifetime. In order to both self-renew and also generate daughter cells that ultimately differentiate into sperm, the SSC pool must undergo repeated rounds of DNA replication.

The process of DNA replication poses a challenge to the fidelity of the genome due to both errors introduced by the DNA polymerases as well as impediments encountered by the polymerases (Mcculloch and Kunkel, 2008). Chromatin also poses additional challenges for eukaryotic cells as during DNA replication nucleosomes must be disassembled ahead of the replication fork and re-allocated in the newly synthesised daughter strands (Ransom et al., 2010). To do this, eukaryotic cells evolved a set of replication-associated histones chaperones. The CAF-1 (Chromatin Assembly Factor-1) complex and ASF1 (Anti-Silencing Function 1) are among the best characterised replication-associated histone chaperones and are recruited through interactions with key replisome components (Franco et al., 2005; Groth et al., 2007; Shibahara and Stillman, 1999). Demonstrating the importance of histone chaperones for DNA replication and cell cycle progression, loss of CAF-1 or ASF1 results in impaired S-phase progression and DNA synthesis (Groth et al., 2005; Hoek and Stillman, 2003; Sanematsu et al., 2006; Schulz and Tyler, 2006).

Given the highly proliferative capacity of germ cells and dramatic changes to chromatin encountered during their development, it is likely that unique challenges must be overcome during gametogenesis. Recent reports support this idea showing that histone chaperones are critical at distinct steps of gametogenesis. For example, the homolog of the P150 large subunit of CAF1 is essential for the maintenance of gonadal stem cells in female *Drosophila* (Clémot et al., 2018). In mice, a homolog of ASF1, ASF1B, supports initiation of meiosis (Messiaen et al., 2014). ASF1B is so far the only histone chaperone identified to be important for gametogenesis in mammals. Generally, the chromatin transactions during DNA replication in mammalian SSCs are poorly characterised. Given the importance of histone chaperone activity for DNA replication and the proliferative nature of SSCs, it is likely that male fertility will be particularly reliant on histone chaperones.

Germ Cell Nuclear Acidic protein (GCNA) has been extensively used as a germ cell marker for almost thirty years (Enders and May, 1994; Tanaka et al., 1997). Recently, mutations in *GCNA* have been linked to azoospermia in humans, defining GCNA as a clinical determinant of human infertility (Arafat et al., 2021; Hardy et al., 2021). Despite this, little is known about the function in mammals of this evolutionarily conserved factor. GCNA contains a SprT protease domain closely related to that found in SPRTN, a critical factor in DNA-protein crosslink (DPC) repair (Carmell et al., 2016). Studies in invertebrates report a role for GCNA in DPC repair in the germline (Bhargava et al., 2020; Borgermann et al., 2019; Dokshin et al., 2020). However, mouse GCNA lacks the SprT protease domain suggesting a role independent of DPC repair. Consistent with this, we find that GCNA is dispensable for maintaining cellular resistance to DPC-causing agents in mice. Instead, we observe that both mouse GCNA and human GCNA interact with core histones and that mouse GCNA (mGCNA) has histone chaperone activity. Moreover, we find that mGCNA can associate with the DNA replication machinery and consistent with a role as a histone chaperone supporting DNA replication, GCNA-deficient SSCs accumulate in S-phase. Consequently, GCNA deficient mice fail to maintain the SSC pool through their lifetime, leading to an age-dependent reduction in sperm production. Therefore, our results suggest that GCNA is a histone chaperone necessary for maintenance of SSCs in mammals.

## Results

### GCNA supports long term gametogenesis in male mice

Although mutations in the *GCNA* locus have been associated with azoospermia in humans, the mechanism behind this is not understood (Arafat et al., 2021; Hardy et al., 2021). Therefore, we sought to define the aetiology of this defect using a previously generated *Gcna* knock-out mouse allele (Appendix Fig S1, Δ Exon 4 allele)(Carmell et al., 2016) (MGI ID: 5910931) JAX (stock ID: 031055). The allele was first validated by western blot and immunofluorescence using two well characterised antibodies (Tra98 and GCNA-1) that recognise the C-terminus of mouse GCNA, a region outside the deletion (Carmell et al., 2016; Enders and May, 1994; Tanaka et al., 1997). We were unable to detect a GCNA signal in the knock-out mice, confirming loss of the protein (Fig 1A, B and C).

**Figure 1 -.**
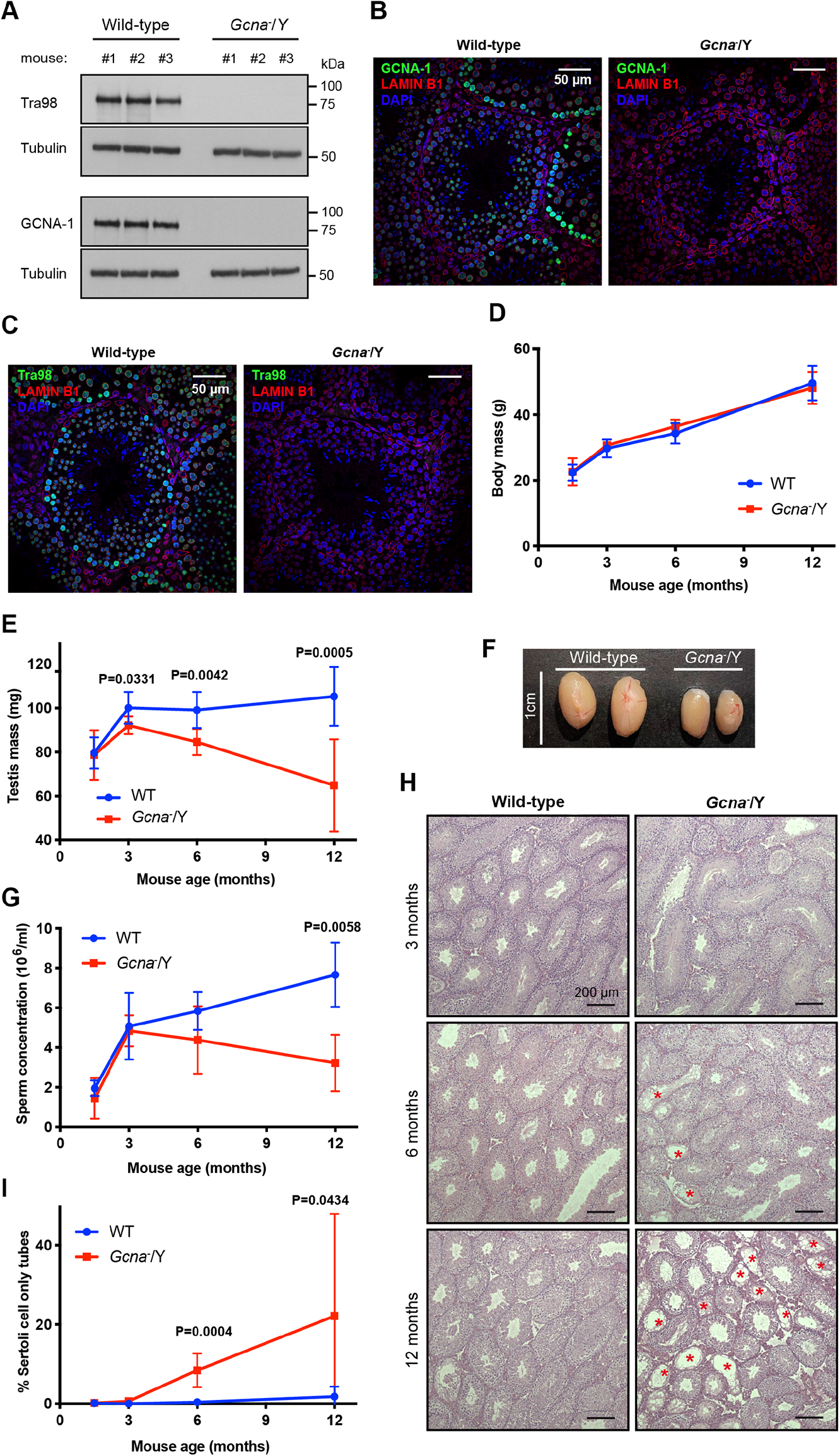
GCNA-deficient male mice exhibit an age dependent decrease of gametes production. A. Validation of *Gcna*^−^/Y mice by Western blot and by using mouse testis lysates. Blots were probed with two antibodies directed against mouse GCNA (Tra98 and GCNA-1) and with an anti-tubulin antibody. Data is representative from two independent experiments. B. Validation of *Gcna*^−^/Y mice by immunofluorescence of testis sections, using the GCNA-1 antibody. LAMIN B1 was displayed in red, GCNA-1 in green and DNA (DAPI stained) in blue. C. Validation of *Gcna*^−^/Y mice by immunofluorescence of testis sections, using the Tra98 antibody. LAMIN B1 was displayed in red, Tra98 in green and DNA (DAPI stained) in blue. D. Body weight of wild-type and *Gcna*^−^/Y male mice between 1.5 months and 12 months. 1.5 months WT (n=6 mice) and *Gcna*^−^/Y (n=5 mice), 3 months WT (n=8 mice) and *Gcna*^−^/Y (n=6 mice), 6 months WT (n=7 mice) and *Gcna*^−^/Y (n=6 mice), 12 months WT (n=9 mice) and *Gcna*^−^/Y (n=6 mice). Data represent the mean and S.D.. E. Testis weight of wild-type and *Gcna*^−^/Y male mice between 1.5 months and 12 months. 1.5 months WT (n=6 mice) and *Gcna^−^*/Y (n=5 mice), 3 months WT (n=8 mice) and *Gcna^−^*/Y (n=6 mice), 6 months WT (n=7 mice) and *Gcna*^−^/Y (n=6 mice), 12 months WT (n=9 mice) and *Gcna^−^/*Y (n=6 mice). Data represent the mean and S.D.. *P* values were calculated by using an unpaired t-test. F. Photograph of representative testes of 1 year old wild-type and *Gcna*^−^/Y mice. G. Sperm concentration obtained per epididymis of wild-type and *Gcna*^−^/Y male mice between 1.5 months and 12 months. 1.5 months WT (n=6 epididymes) and *Gcna*^−^/Y (n=5 epididymes), 3 months WT (n=8 epididymes) and *Gcna*^−^/Y (n=6 epididymes), 6 months WT (n=6 epididymes) and *Gcna*^−^/Y (n=6 epididymes), 12 months WT (n=8 epididymes) and *Gcna*^−^/Y (n=5 epididymes). Data represent the mean and S.D.. *P* value was calculated by using an unpaired t-test. H. Micrographs of Haematoxylin and Eosin stained testis sections of 3, 6 and 12 months old wild-type and *Gcna^−^*/Y mice. Red asterisks highlight Sertoli Cells-Only seminiferous tubes (SCOs). I. Frequency of Sertoli Cells-Only seminiferous tubes in testes of wild-type and *Gcna^−^*/Y male mice between 1.5 months and 12 months. A minimum of 70 seminiferous tubes are scored per mouse. 1.5 months WT (n=6 mice) and *Gcna*^−^/Y (n=5 mice), 3 months WT (n=8 mice) and *Gcna*^−^/Y (n=6 mice), 6 months WT (n=7 mice) and *Gcna*^−^/Y (n=6 mice), 12 months WT (n=8 mice) and *Gcna^−^*/Y (n=5 mice). Data represent the mean and S.D.. *P* values were calculated by using an unpaired t-test.

We first set out to ask if GCNA was required for mouse development. GCNA-deficient male mice were born at Mendelian ratios (Fig EV1A and B). Furthermore, the body weights and one year survival of male mice lacking GCNA were indistinguishable from wild-type littermates (Figs 1D and EV1C and D). We then confirmed that the expression of GCNA is restricted to the testis, in adult mice (Fig EV1E)(Enders and May, 1994; Tanaka et al., 1997). As these results suggest a specific role of GCNA in testis physiology, we then focused our attention on the testis development of *Gcna* knock-out mice. Testis mass at 6 weeks old is indistinguishable from wild-type males (Fig 1E). However, it decreases over time in the absence of GCNA, but not in wild-type controls. By one year of age, GCNA deficient males show approximately a 40% reduction in testis mass compared to littermate controls (Fig 1E and F). Consistently, the testis weight reduction is accompanied by a time-dependent decrease of sperm number in the epididymis (Fig 1G). We did not observe any difference in the mass of the seminal vesicle nor the serum testosterone concentration suggesting that hormonal signalling is intact in the absence of GCNA and it is not causing the reduction of the sperm production (Fig EV1F and G).

In contrast to the previous report with mice carrying the same *Gcna* allele, we did not observe a complete loss of sperm in the epididymis as described previously (Fig EV1H)(Carmell et al., 2016). This discrepancy may in part be explained by the use of different Cre recombinases, *Mvh*^Cre-mOrange^ in the previous study and *Stella-Cre* in this study, to convert the floxed allele to a knockout (Hu et al., 2013)(MGI ID: 6505285) JAX (stock ID: 035877) (Liu et al., 2011)(MGI ID: 5004882). Following recombination, we backcrossed the recombined *Gcna* allele onto a C57 background ensuring that we used mice carrying a germline GCNA mutant allele but not Cre-recombinase for subsequent breeding. Then, we went on to show that we were studying a null allele of *Gcna*. Firstly, the recombined allele resulted in excision of the floxed exon (Appendix Fig S1B). Secondly, mice carrying the recombined allele did not express the GCNA protein, when assayed by immunofluoresence and immunoblotting (Fig 1A-C). This provides strong evidence that the allele of *Gcna* used in our study does not produce functional protein. Although *Gcna* knock-out male mice in our study produce less sperm upon ageing, these remaining sperm are functional (Fig EV1I). Together, our observations reveal that whilst spermatogenesis is not blocked in the absence of GCNA, males lacking GCNA are unable to sustain sperm production over time.

To determine the basis of this defect, we histologically assessed the testes at different ages (Fig 1H). We observed a time-dependent increase in seminiferous tubes lacking germ cells, or Sertoli cell only tubes (SCOs), in *Gcna* knock-out male mice (Fig 1I). These results provide an explanation for the decrease of the testis mass and sperm production over time, showing that GCNA supports the maintenance of the male germ line with time.

As GCNA is important for homeostasis of male germ cells, we sought to assess if it also has a role in females. *Gcna* knock-out female mice were born at the Mendelian ratio, suggesting that GCNA is not necessary for female development (Fig EV2A and B). We then counted follicles in ovaries of young and aged mice and did not observe any difference between wild-type and *Gcna* knock-out females (Fig EV2C and D). Consistent with this result, no reduction in fertility was observed in *Gcna* knock-out female mice (Fig EV2E, F and G). These results indicate that GCNA is dispensable for the female reproductive function and that maternal GCNA is dispensable for embryonic development.

### GCNA preserves the spermatogonial stem cell pool in mice

We next sought to identify the cause of the age-dependent decrease in spermatogenesis in mice lacking GCNA. We first tested if the premature loss of germ cells could be due to a seeding defect of the embryonic gonads. We therefore crossed the *Gcna* knock-out allele with a primordial germ cell reporter which expresses GFP under the control of a fragment of the *Oct4* promoter (also known as GOF18-GFP)(Szabo et al., 2002)(Appendix Fig S2A). We found that E12.5 *Gcna* knock-out males have indistinguishable numbers of PGCs compared to wild-type littermates (SSEA1^+^ GOF18-GFP^+^ PGGs, Appendix Fig S2B, C and D). These results suggest that the age-dependent loss of germ cells in males lacking GCNA may not have an embryonic origin.

Our results indicate that GCNA-deficient male mice have normal numbers of germ cells and normal testis mass and sperm production in their first weeks of life. We hypothesised therefore that the spermatogonial stem cell (SSC) pool prematurely contracts during the post-natal life, in the absence of GCNA. To test this, we first verified that GCNA is expressed in SSCs by immunofluorescence. SSCs were identified by immunostaining of the SSC marker promyelocytic leukaemia zinc finger (PLZF) and we observed that a high proportion of PLZF positive cells (approximatively 70%) also expressed GCNA (Fig 2A and B). Consistent with a role for GCNA in human spermatogenesis too, we observed the same expression pattern in human SSCs (Fig 2C and D).

**Figure 2 -.**
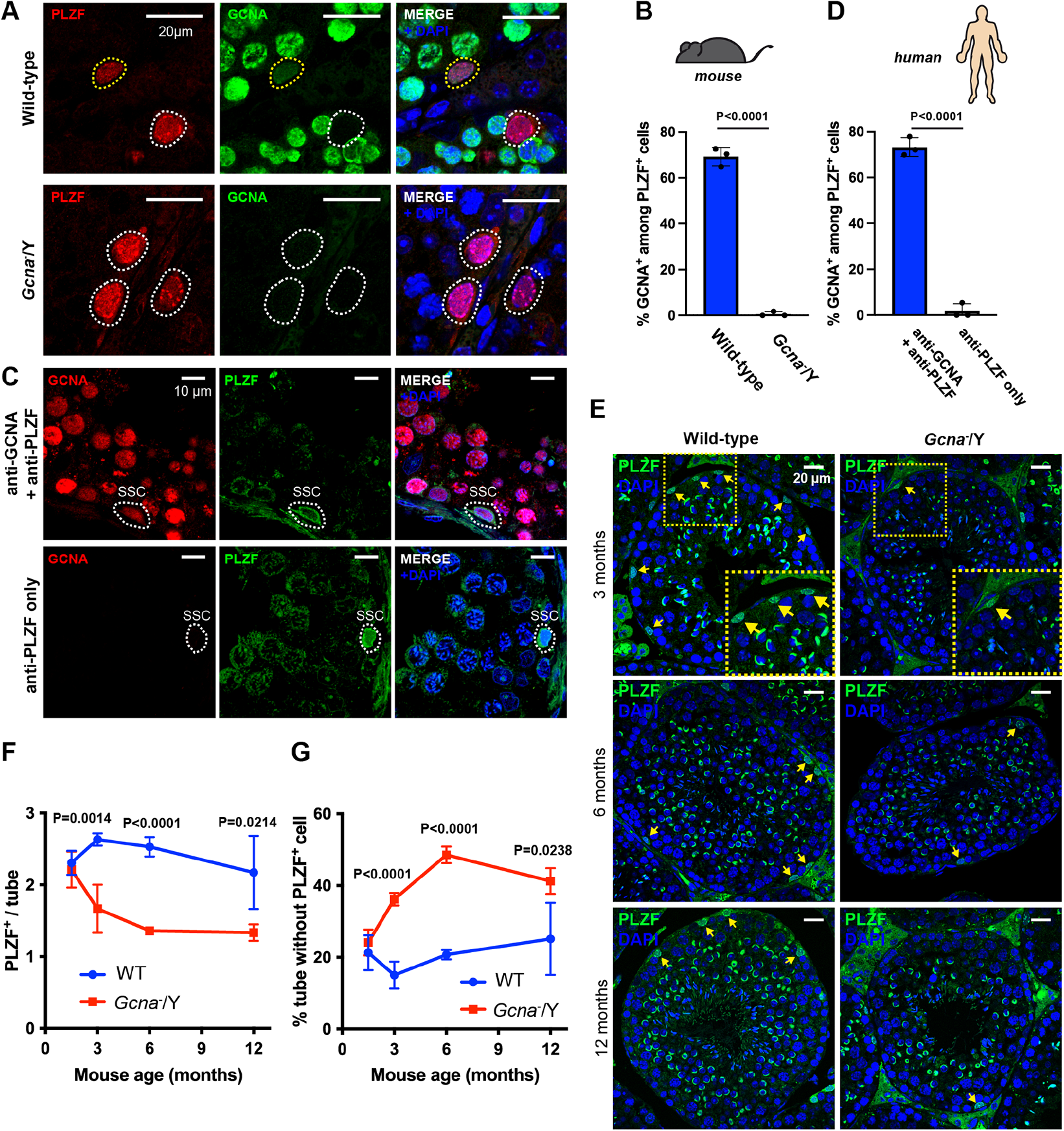
Age dependent loss of spermatogonial stem cells in absence of GCNA. A. Immunofluorescence staining of GCNA (Tra98) and PLZF on adult mice testis sections. PLZF was displayed in red, GCNA in green and DNA (DAPI stained) in blue. Cells positive for both PLZF and GCNA are highlighted in yellow and PLZF only cells are highlighted in white. GCNA is expressed in mice SSCs. B. Frequency of PLZF positive cells also positive for GCNA in mice. A minimum of 50 PLZF positive cells are scored per mouse. Data represent the mean and S.D.. N=3 mice for each genotype. *P* value was calculated by using an unpaired t-test. C. Immunofluorescence staining of GCNA and PLZF on human testis sections. GCNA was displayed in red, PLZF in green and DNA (DAPI stained) in blue. GCNA is expressed in human SSCs. D. Frequency of PLZF positive cells also positive for GCNA in human. A minimum of 50 PLZF positive cells are scored per experiment. Data represent the mean and S.D.. Data represent three independent experiments. *P* value was calculated by using an unpaired t-test. E. Immunofluorescence staining of PLZF on testis sections of 3, 6 and 12 months old wild-type and *Gcna*^−^/Y mice. PLZF is displayed in green and the DNA staining DAPI in blue. Yellow arrows highlight SSCs, which are PLZF positive. Bottom panels are magnifications of regions highlighted by the yellow boxes. F. Number of PLZF positive cells per seminiferous tube in wild-type and *Gcna*^−^/Y testes, between 1.5 months and 12 months. A minimum of 50 seminiferous tubes are scored per mouse. Data represent the mean and S.D.. N=4 mice for each genotype and age. *P* values were calculated by using an unpaired t-test. G. Frequency of seminiferous tubes displaying no PLZF positive cells in wild-type and *Gcna*^−^/Y testes, between 1.5 months and 12 months. A minimum of 50 seminiferous tubes are scored per mouse. Data represent the mean and S.D.. N=4 mice for each genotype and age. *P* values were calculated by using an unpaired t-test.

We then tested if GCNA is important for the maintenance of the SSC population in mice. The SSC population in adult animals (10 weeks old) was first assessed by quantification of PLZF by western blot. Interestingly, we observed reduced PLZF expression in the GCNA-deficient testes (Fig EV3A andB). In contrast, we did not observe any reduction of the expression of the Sertoli cell marker Wilms tumor 1 (WT1) suggesting that the germ cell niche is not altered in *Gcna* knock-out male mice (Fig EV3A and C). These results suggest a reduction in the number of SSCs or a reduced expression of PLZF.

We therefore assessed the number of SSCs in GCNA-deficient male mice and how this number changes over mice lifetime. We counted PLZF positive SSCs in tubes exhibiting spermatogenesis, excluding SCO tubes, at different ages. As observed with the testis mass and the sperm concentration, a time-dependent decrease in the SSC number was found, first apparent at 3 months of age (Fig 2E and F). Consistent with this result, there was an age-dependent increase in the proportion of seminiferous tubes lacking any PLZF positive SSC (Fig 2G). These data suggest that GCNA is required to maintain the SSC population over time in mice, which permits the continuous production of sperm.

### Neither apoptosis nor premature differentiation drive age-dependent loss of SSCs in *Gcna* knock-out males

As SSCs in GCNA-deficient male mice are lost in an age-dependent manner, we sought to elucidate how these cells are lost. As SSCs can either be lost through apoptosis or differentiation, we first tested these two fates (Diao et al., 2016; Fu et al., 2018). To test if apoptosis was increased in SSCs, we measured the frequency of cleaved-caspase 3 (CC3) positive SSCs (Fig EV4A and B). We could not detect any CC3 positive SSC in either wild-type or GCNA-deficient testes, at 1.5 or 6 months old. Furthermore, we could not detect any difference in the frequency of apoptotic germ cells between either wild-type and GCNA-deficient testes (Fig EV4C, D and E). We then tested a potential clearance of defective SSCs through increased differentiation rate by assessing the frequency of SSCs expressing DNMT3B, a factor that drives differentiation of SSCs (Shirakawa et al., 2013). Frequencies of differentiating SSCs were similar between wild-type and GCNA-deficient animals (Fig EV4F and G). Together, these results suggest that neither increased apoptosis nor differentiation of SSCs is the origin of the age-dependent loss of SSCs in *Gcna* knock-out males.

### GCNA is necessary for S-phase normal progression in murine SSCs

As apoptosis or differentiation do not explain the SSC loss in absence of GCNA, we tested an alternative hypothesis. It has been shown that reduced quiescence triggers premature exhaustion of the SSC pool (Liao et al., 2014; Takubo et al., 2007). To test if this mechanism could explain the age-dependent loss of SSCs in *Gcna* knock-out males, we assessed the frequency of SSCs negative for the proliferation marker Ki67 (Fig 3A). Interestingly, whilst the proportion of quiescent SSCs increases with age in wild-type mice, no such increase is observed in the absence of GCNA (Fig 3B). These results suggest that GCNA-deficient SSCs remain more frequently in the cell cycle, unlike wild-type SSCs.

**Figure 3 -.**
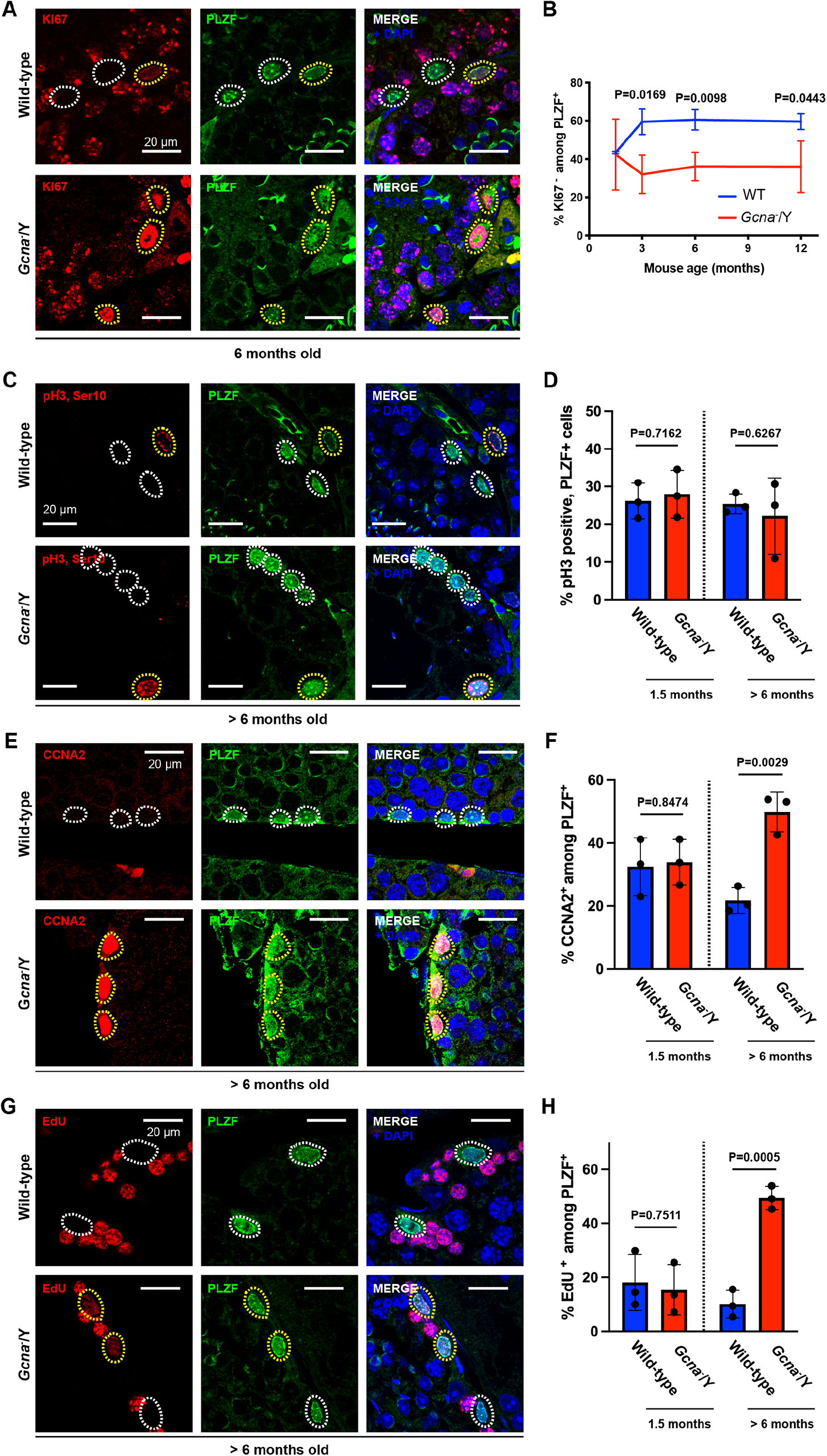
Accumulation of spermatogonial stem cells in S-phase in absence of GCNA. A. Immunofluorescence staining of wild-type and *Gcna*^−^/Y testis sections from 6 months old mice. KI67 was displayed in red, PLZF in green and DNA (DAPI stained) in blue. Cells positive for both PLZF and KI67 are highlighted in yellow and PLZF only cells are highlighted in white. B. Frequency of PLZF positive cells negative for KI67 (quiescent SSCs) at 1.5, 3, 6 and 12 months old. A minimum of 50 PLZF positive cells are scored per mouse. Data represent the mean and S.D.. N=3 mice for each genotype and age. *P* values were calculated by using an unpaired t-test. C. Immunofluorescence staining of wild-type and *Gcna*^−^/Y testis sections from >6 months old mice. Phosphorylation of H3 (pH3, Ser10) was displayed in red, PLZF in green and DNA (DAPI stained) in blue. Cells positive for both PLZF and pH3 are highlighted in yellow and PLZF only cells are highlighted in white. D. Frequency of PLZF positive cells also positive for pH3 at 1.5 and >6 months old. A minimum of 50 PLZF positive cells are scored per mouse. Data represent the mean and S.D.. N=3 mice for each genotype and age. *P* values were calculated by using an unpaired t-test. E. Immunofluorescence staining of wild-type and *Gcna*^−^/Y testis sections from >6 months old mice. Cyclin A2 (CCNA2) was displayed in red, PLZF in green and DNA (DAPI stained) in blue. Cells positive for both PLZF and CCNA2 are highlighted in yellow and PLZF only cells are highlighted in white. F. Frequency of PLZF positive cells also positive for CCNA2 at 1.5 and >6 months old. A minimum of 50 PLZF positive cells are scored per mouse. Data represent the mean and S.D.. N=3 mice for each genotype and age. *P* values were calculated by using an unpaired t-test. G. Immunofluorescence staining of wild-type and *Gcna*^−^/Y testis sections from >6 months old mice. EdU was given to mice through intraperitoneal injection and mice were culled for histological analyses 4h after injection. EdU was displayed in red, PLZF in green and DNA (DAPI stained) in blue. Cells positive for both PLZF and EdU are highlighted in yellow and PLZF only cells are highlighted in white. H. Frequency of PLZF positive cells also positive for EdU at 1.5 and >6 months old. A minimum of 50 PLZF positive cells are scored per mouse. Data represent the mean and S.D.. N=3 mice for each genotype and age. *P* values were calculated by using an unpaired t-test.

As GCNA-deficient SSCs appear committed to the cell cycle, we tested if these cells are also frequently dividing. We therefore assessed the frequency of SSCs positive for the M phase marker H3 pSer10. Surprisingly, old *Gcna* knock-out male mice (> 6 months old) do not exhibit an increased frequency of H3 pSer10 positive SSCs compared to wild-type, indicating that GCNA-deficient SSCs are not dividing more frequently (Fig 3C and D). The observation that more SSCs are engaged in the cell cycle but are not dividing more suggests that GCNA-deficient SSCs may be accumulating in a particular phase of the cell cycle.

We consequently tested if SSCs accumulate in S/G2, by using the cyclin A2 (CCNA2) marker. Interestingly, we found a significant increase in CCNA2 positive SSCs, in old GCNA-deficient male mice suggesting an accumulation in S/G2 (Fig 3E and F). We then decided to narrow down the SSC defect by looking more closely at S-phase using the incorporation of the nucleotide analogue named EdU. Strikingly, we observed a significant increase in the frequency of EdU positive SSCs in old GCNA-deficient male mice suggesting these SSCs are more frequently in S-phase than wild-type SSCs (Fig 3G and H). Together with the previous data on the cell cycle properties of GCNA-deficient SSCs, this result suggests that there is likely an elongation of S-phase in SSCs of old mice lacking GCNA.

### Mouse GCNA associates with the replication machinery

Given our observations, we sought to understand how GCNA can support the progression of S-phase. Therefore, we tested if GCNA associates with chromatin, specifically during S-phase. Subcellular fractionation experiments showed that endogenous mouse GCNA (mGCNA) was observed in the chromatin-associated fraction of asynchronous cultures of mouse embryonic stem cells (mESCs) (Fig 4A). Additionally, we observed that this association with chromatin was sensitive to DNAse treatment. We went on to show by immunofluorescence in 3T3 cells ectopically expressing mGCNA, that mGCNA remains nuclear after pre-extraction and this is DNA-dependent too (Fig 4B). We then focused our attention on cells in S-phase. Cell fractionation of synchronised cells showed that mGCNA remains in the chromatin-associated fraction of S-phase cells (Fig 4C). We also confirmed by immunofluorescence that mGCNA remains in the chromatin of replicating cells (EdU positive)(Fig 4D and E). Taken together, these experiments strongly suggest that mGCNA is a chromatin associated protein and this association remains in S-phase.

**Figure 4 -.**
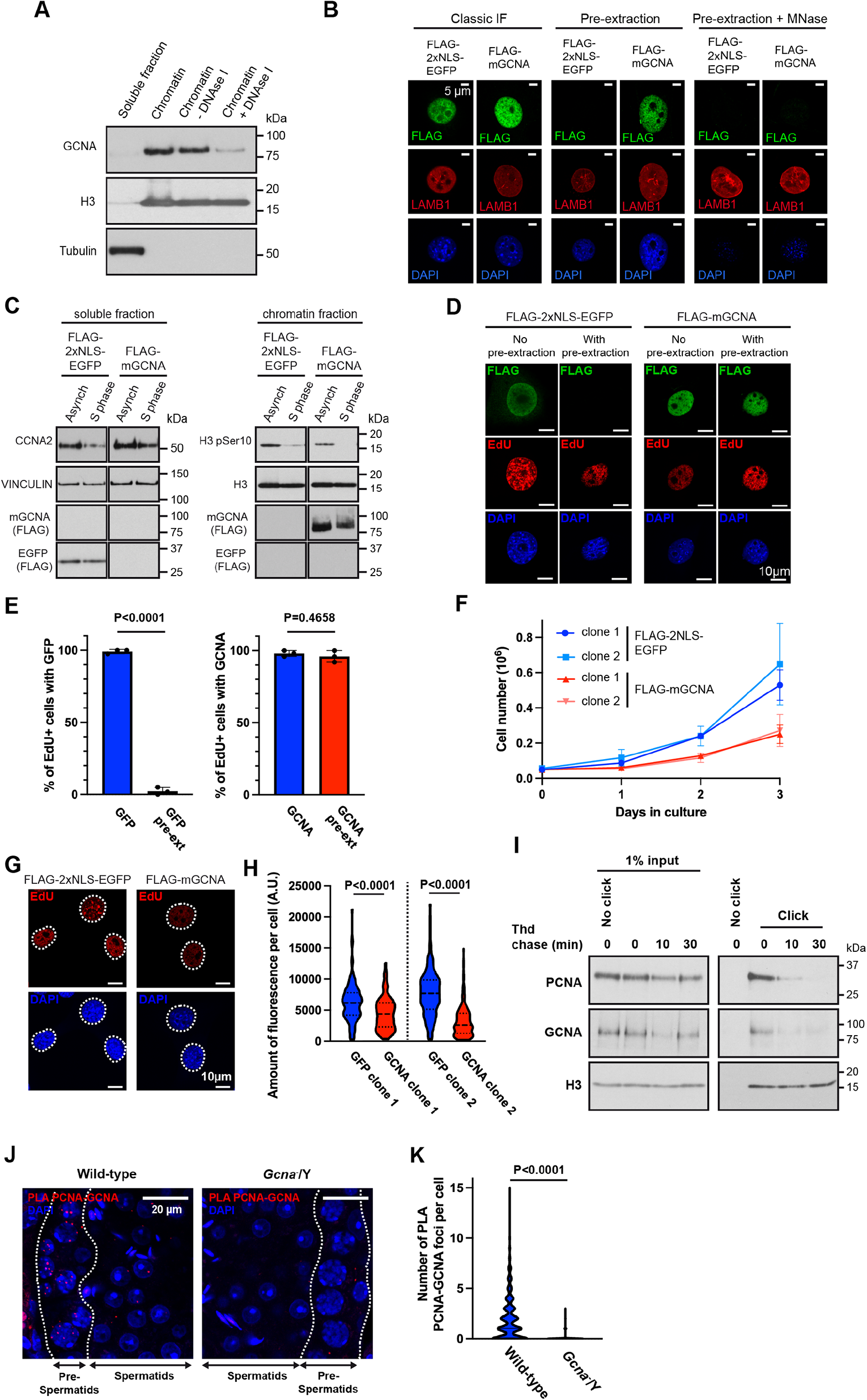
Association of mGCNA with the DNA replication machinery. A. Western blot of cell fractionations performed on BAC9 mESC. Chromatin fraction was incubated or not with DNAse I to digest DNA. The blot was probed with anti-GCNA (GCNA-1), anti-H3 and anti-tubulin antibodies. Data is representative from three independent experiments. B. Immunofluoresence of FLAG-2xNLS-EGFP or FLAG-mGCNA in stably transfected 3T3 cells. 3T3 cells are stably expressing FLAG-2xNLS-EGFP or FLAG-mGCNA. Cells were fixed then permeabilised (classic IF) to access the whole fraction of the proteins or pre-permeabilised then fixed (pre-extraction) to access the chromatin fraction. Data is representative from three independent experiments. C. Western blots of cell fractionations performed on asynchrone or S-phase synchronised 3T3 cells. 3T3 cells are stably expressing FLAG-2xNLS-EGFP or FLAG-mGCNA. Blots were probed with anti-Cyclin A2 (CCNA2), anti-VINCULIN, anti-FLAG, anti-H3 phospho Ser10 and anti-H3. Data is representative from three independent experiments. D. Immunofluoresence of FLAG-2xNLS-EGFP or FLAG-mGCNA in stably transfected 3T3 cells. Prior fixation or permeabilisation, cells were incubated with 10µM EdU for 30 min. Cells were fixed then permeabilised to access the whole fraction of the proteins (No pre-extraction) or pre-permeabilised then fixed to access the chromatin fraction (pre-extraction). Data is representative from three independent experiments. E. Frequency of EdU positive cells exhibiting nuclear EGFP or mGCNA without or with prextraction, as displayed in (D). Data represent the mean and S.D.. Data represent three independent experiments. *P* values were calculated by using an unpaired t-test. F. Growth curves of 3T3 clones stably expressing FLAG-2xNLS-EGFP or FLAG-mGCNA. Data represent the mean and S.D.. Data represent three independent experiments. G. Micropgraphs of the labelling of EdU, after a 30 min pulse at 10µM, in 3T3 cells stably expressing FLAG-2xNLS-EGFP or FLAG-mGCNA. H. Quantification of the amount of fluorescence (integrated density) of EdU per cell, in different 3T3 clones stably expressing FLAG-2xNLS-EGFP or FLAG-mGCNA. Clones were processed and analysed pair-wise. FLAG-2xNLS-EGFP clone 1 (N=210 cells), FLAG-mGCNA clone 1 (N=185 cells), FLAG-2xNLS-EGFP clone 2 (N=213 cells), FLAG-mGCNA clone 2 (N=204 cells). Data represent the median and interquartile range. *P* values were calculated by using a two-tailed Mann-Whitney test. I. Immunoblot of proteins precipitated during the iPOND from BAC9 mESC. Cells were pulsed with EdU for 10 min and then incubated with thymidine (Thd) for 0, 10 and 30 min. iPOND was then performed and eluted proteins were analysed by Western blot. The blot was probed with anti-PCNA, anti-GCNA (Tra98) and anti-H3 antibodies. Without click chemistry (No click lane), no proteins are precipitated from nascent DNA. PCNA and GCNA are present at the nascent DNA (Click, t0 min) but not on DNA after the thymidine chase (Click, t10 and 30 min). Data is representative from two independent experiments. J. Proximity ligation assay performed with anti-PCNA and anti-GCNA (Tra98) antibodies on wild-type and *Gcna*^−^/Y adult testis sections. PLA foci are abundant in pre-spermatid cells, were PCNA and GCNA are co-expressed. Data is representative from three independent experiments. K. Quantification of the number of PLA PCNA-GCNA foci per pre-spermatid cells. Wild-type (N=1892 cells, in 3 mice), *Gcna*^−^/Y (N=1919 cells, in 3 mice). Data represent the mean and S.D.. *P* value was calculated by using a two-tailed Mann-Whitney test.

Our experiments show that the association between mGCNA and chromatin is sensitive to DNAse treatment. We therefore tested if mGCNA could bind DNA by DNA pulldown. Mouse SPRTN was used as a positive control as it is a paralog of GCNA and it has been shown to bind single- and double-stranded DNA (ssDNA and dsDNA)(Stingele et al., 2016). We used a catalytically inactive SPRTN (E113Q) as binding of SPRTN with DNA activates its protease activity and triggers autocleavage of the protein, making SPRTN detection difficult. We did not observe any interaction between mGCNA and DNA, suggesting that mGCNA could be recruited to chromatin through an interaction with a partner (Appendix Fig S3).

As GCNA is associated with chromatin in S-phase but does not directly bind to DNA, we sought to investigate a functional interaction of mGCNA with the DNA replication. We first observed that expression of mGCNA reduces the proliferation of 3T3 cells (Fig 4F). Then, we set out to test if this reduced proliferation could be due to mGCNA interfering with S-phase progression. We therefore investigated the DNA replication rate of 3T3 cells by measuring the incorporation of EdU, in replicating cells. Microscopic analysis showed that ectopic mGCNA expression reduces the incorporation of EdU (Fig 4G, H). This result suggests that mGCNA interferes with DNA replication when it is expressed in cells.

Given the S-phase association of mGCNA with chromatin, its interference with replication but lack of direct DNA binding, we sought to directly test if there was an association of mGCNA with the DNA replication machinery. To answer this question, we performed isolation of proteins on nascent DNA in mESCs (iPOND)(Sirbu et al., 2012). Ubiquitously chromatin-bound proteins, such as H3, remain detectable both at the replication fork, i.e. after EdU pulse, and in thymidine-chased samples (Fig 4I). However, replication-associated proteins like PCNA are enriched on the nascent DNA, after the EdU pulse. Interestingly, we observed that GCNA associates with nascent DNA, as does PCNA. Furthermore, the enrichment of both proteins is gradually lost from the nascent DNA during the thymidine chase. These results suggest that, as PCNA, GCNA can associate with the DNA replication fork.

We then sought to assess if GCNA associates with DNA synthesis in testes, where we have shown a physiological role of GCNA. To test this, we analysed its co-localisation with PCNA and observed a high frequency of germ cells co-expressing both PCNA and GCNA in adult testes (Appendix Fig S4). Cells co-expressing both factors are predominantly in the pre-spermatid stages of the spermatogenesis so we sought to test whether GCNA and PCNA are in close proximity in pre-spermatid cells. Consistent with the co-expression pattern, we observed by proximity ligation assay (PLA) that pre-spermatid cells exhibited specific signals for the proximity of PCNA and GCNA (Fig 4J and K). These results suggest that PCNA and GCNA associate in pre-spermatid cells, potentially during DNA replication and DNA synthesis.

### GCNA is dispensable for DPC repair in mice

In invertebrates, GCNA is required for the repair of DNA-protein crosslinks (DPCs) during embryonic development and in the germ line (Bhargava et al., 2020; Borgermann et al., 2019; Dokshin et al., 2020). DPCs impede S-phase progression by inducing replication fork collapse, eventually leading to formation of DNA breaks (Stingele et al., 2017). Here we observe that mGCNA associates with the DNA replication, a localisation compatible with a role in DPC repair. Therefore, we tested if the absence of GCNA leads to DNA breaks formation in SSCs, explaining their accumulation in S-phase. To do this, we assessed the presence of DNA breaks in SSCs by using the γH2AX marker. We did not observe any difference between wild-type and GCNA-deficient male mice in the frequency of SSCs positive for γH2AX nor in the γH2AX signal intensity in SSCs, at 1.5 or 6 months old (Fig 5A, B and C and Appendix Fig S5). These results suggest that GCNA-deficient SSCs are not likely accumulating in S-phase because of DNA breaks.

**Figure 5 -.**
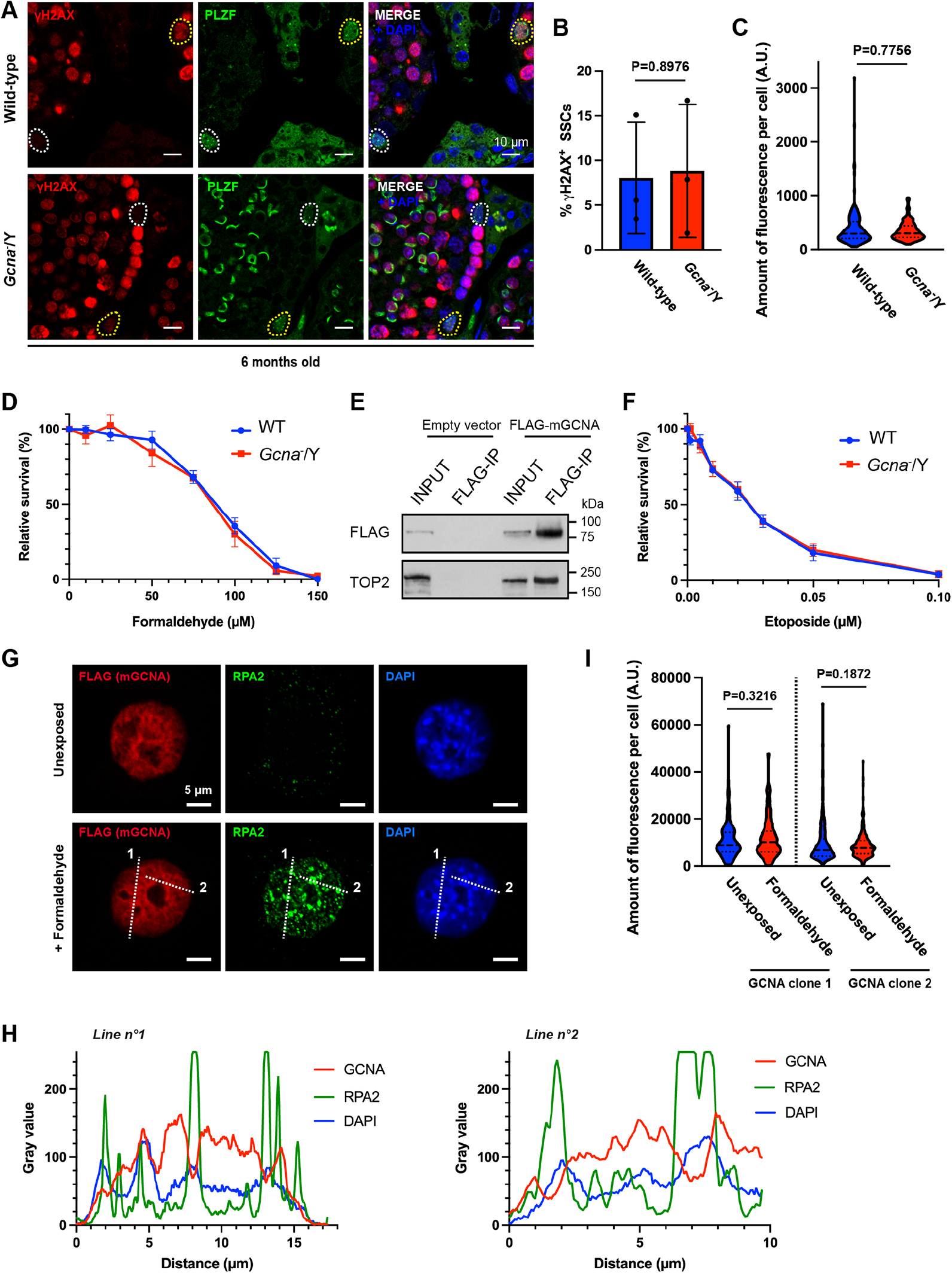
Non-requirement of GCNA for DPC repair in mice. A. Immunofluorescence staining of wild-type and *Gcna*^−^/Y testis sections from 6 months old mice. γH2AX was displayed in red, PLZF in green and DNA (DAPI stained) in blue. Cells positive for both PLZF and γH2AX are highlighted in yellow and PLZF only cells are highlighted in white. B. Frequency of PLZF positive cells also positive for γH2AX at 6 months old. A minimum of 50 PLZF positive cells are scored per mouse. Data represent the mean and S.D.. N=3 mice for each genotype. *P* values were calculated by using an unpaired t-test. C. Quantification of the amount of fluorescence (integrated density) of γH2AX per SSC, in wild-type and *Gcna*^−^/Y testis from 6 months old mice. Wild-type (N=161 SSCs from three mice), *Gcna*^−^/Y (N=151 SSCs from three mice). Data represent the median and interquartile range. *P* values were calculated by using a two-tailed Mann-Whitney test. D. *Gcna*^−^/Y mESC are not sensitive to formaldehyde. Data represent the mean and S.D.; data represent three independent experiments each carried out in triplicate. E. Western blot of the FLAG immunoprecipitation. 3T3 cells were transiently transfected with FLAG-mGCNA and a FLAG immunoprecipitation was performed on the soluble fraction. The blot was probed with anti-FLAG and anti-TOP2 (recognising both TOP2α and TOP2β) antibodies. Data is representative from three independent experiments. F. *Gcna*^−^/Y mESC are not sensitive to etoposide. Data represent the mean and S.D.; data represent three independent experiments each carried out in triplicate. G. Immunofluoresence of 3T3 cells stably expressing FLAG-mGCNA. Cells were first exposed to 600 µM formaldehyde for 1h, in order to induce DPCs. Then, cells were pre-permeabilised and fixed (pre-extraction) to access the chromatin fraction. FLAG (mGCNA) was displayed in red, RPA2 in green and DNA (DAPI stained) in blue. The dotted white lines (1 and 2) highlight the region analysed in (H). H. Profiles of the intensity of signals for GCNA, RPA2 and DAPI along the lines highlighted in (G). I. Quantification of the amount of fluorescence (integrated density) of GCNA (FLAG) per cell, in different 3T3 clones stably expressing FLAG-mGCNA. FLAG-mGCNA clone 1, unexposed (N=206 cells), FLAG-mGCNA clone 1, + formaldehyde (N=211 cells), FLAG-mGCNA clone 2, unexposed (N=216 cells), FLAG-mGCNA clone 2, + formaldehyde (N=211 cells). Data represent the median and interquartile range. *P* values were calculated by using a two-tailed Mann-Whitney test.

Given our results, we sought to directly test if GCNA is involved in DPC repair in mice. To assess the putative role of GCNA in DPC repair in mouse cells, we took advantage of the fact that mESCs express GCNA. Loss of GCNA sensitises *C. elegans* to formaldehyde, which crosslinks a broad spectrum of proteins to DNA (Borgermann et al., 2019). Therefore, we derived mESCs from wild-type or *Gcna^−^*/Y embryos and exposed them to formaldehyde (Appendix Fig S6 and Fig 5D). No significant difference in cellular formaldehyde resistance was observed between GCNA-deficient and wild-type mESCs.

We then asked if mGCNA could be involved in the repair of a specific and frequently formed class of DPC. It has been previously shown that mGCNA interacts with Topoisomerase 2 (TOP2) (Dokshin et al., 2020) suggesting a role in the repair of DNA crosslinked to TOP2. We first confirmed by immunoprecipitation that mGCNA interacts with TOP2 (Fig 5E). We then exposed mESCs to etoposide, which induces DNA-TOP2 crosslinks, and assessed their sensitivity. Again, no significant difference was observed between GCNA-deficient and wild-type mESCs (Fig 5F). These results suggest that mGCNA is dispensable for maintaining cellular resistance to two archetypal DPC-inducing agents, in contrast to the reported role of GCNA in invertebrates.

Human GCNA can form nuclear foci after formaldehyde exposure in U2OS cells (Borgermann et al., 2019). We therefore exposed 3T3 cells stably expressing mGCNA to formaldehyde and assessed its chromatin localisation. We observe that formaldehyde exposure does not change the chromatin localisation of mGCNA and the protein does not form foci (Fig 5G and H). Furthermore, formaldehyde exposure does not alter the association of mGCNA with chromatin in mouse cells (Fig 5I). These data suggest that the chromatin localisation, and role, of mGCNA are DPC-independent.

### Mouse GCNA is closely related to the Intrinsically Disordered Region (IDR) of human GCNA

As mGCNA appears dispensable for DPC repair, we sought to uncover the molecular functions that may be required for SSC self-renewal. To gain insight on this, we first turned to the domain structure of GCNA. GCNA is conserved from *Schizosaccharomyces pombe* to humans and is characterised by the presence of an N-terminal intrinsically disordered region (IDR) and a C-terminal SprT protease domain (Carmell et al., 2016). However, in agreement with the dispensable role in DPC repair, bioinformatic analysis reveals that GCNA in rodents does not possess the SprT protease domain which is essential for DPC repair in invertebrates (Bhargava et al., 2020; Carmell et al., 2016)(Fig EV5A). In order to validate the *in silico* annotation of the mouse gene, we first tested if the endogenous protein in the testis and mESCs had the expected molecular mass of 53 kDa. Using the two characterised antibodies directed against GCNA, i.e. GCNA-1 and Tra98, we observed that the protein migrates between 75 and 100 kDa, consistent with previous reports, but larger than the predicted 53kDa mass (Fig EV5B and C)(Carmell et al., 2016; Enders and May, 1994; Tanaka et al., 1997). Consequently, the cDNA corresponding to the annotated mouse *Gcna* locus was expressed in a heterologous expression system, *E.coli* (Fig EV5B and C). We observed the same migration of the recombinant protein which strongly suggests that the annotation of mGCNA is correct and the protein migration is retarded, likely due to its highly acidic content.

As mGCNA is disordered with highly acidic content, we sought to study the phylogeny of GCNA IDRs among metazoans, particularly focusing our attention on the amount and distribution of charged residues in those IDRs. Interestingly, high content of negatively charged residues is a feature conserved in most mammalian GCNAs (Fig EV5D). Furthermore, GCNA in mammals is characterised by a large acidic region flanked by a basic one (Fig EV5E). This feature was not observed in other metazoan GCNAs examined. The conservation of the acidic region in the IDR of GCNA in mammals, plus the absence of the SprT domain in mouse GCNA suggest that i) the acidic nature of the IDR of GCNA is important for its function and ii) murine GCNA is a separation-of-function model for the study of the IDR of GCNA *in vivo*.

### Mouse and human GCNA bind histones

Our results suggest that mouse GCNA is highly acidic and is a chromatin-associated protein. We therefore hypothesized that the association of mGCNA with chromatin could rely on electrostatic interactions. To test this hypothesis, we washed the mESC chromatin fractions with increasing concentrations of salt and asked if mGCNA remained bound (Fig 6A). Interestingly, mGCNA was almost completely lost at high salt concentrations, supporting an interaction of mGCNA with chromatin through electrostatic interactions.

**Figure 6 -.**
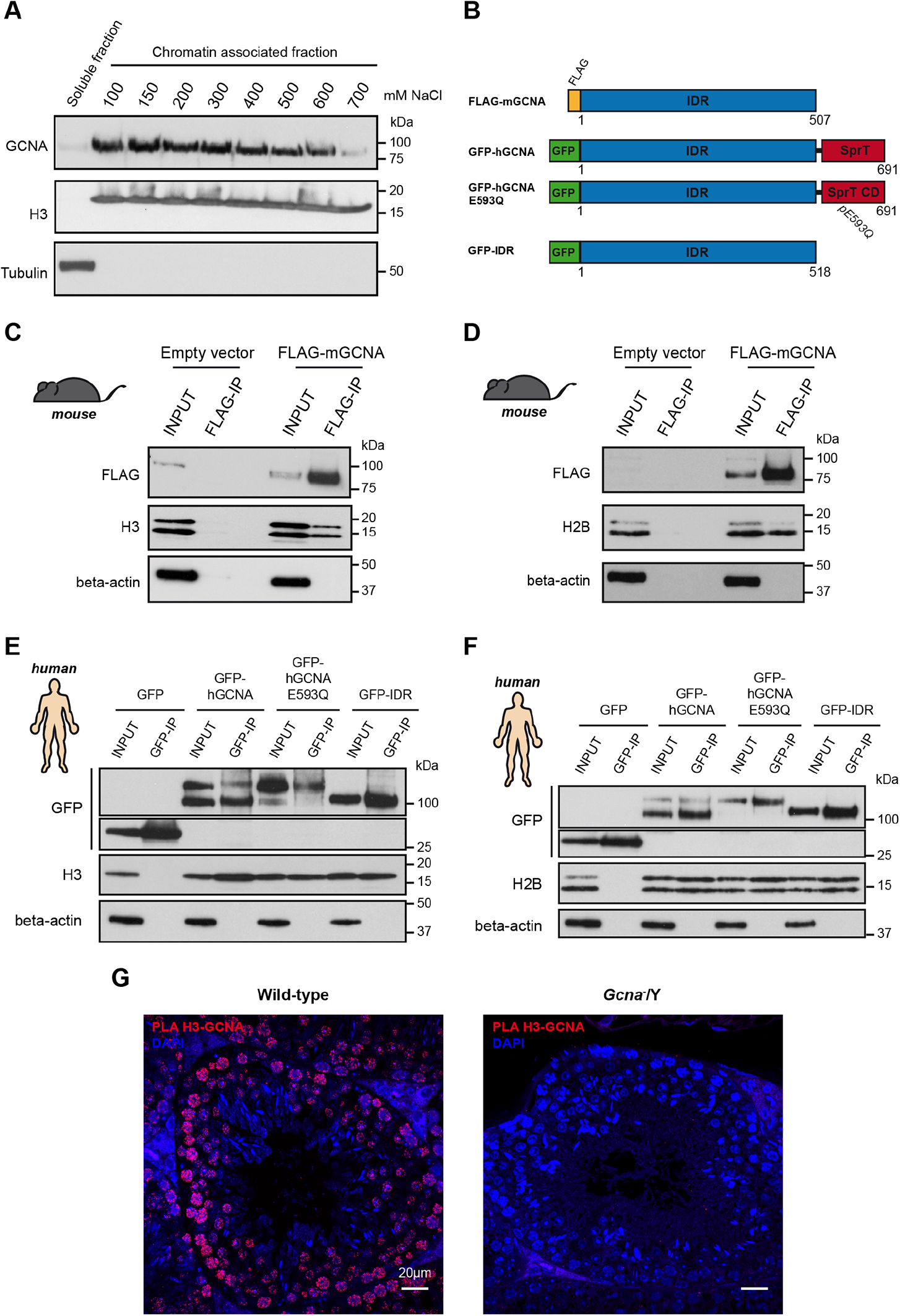
GCNA is a histones binding protein. I. Western blot of cell fractionations performed on BAC9 mESC. Chromatin fraction was washed with increasing concentrations of NaCl to release electrostatic interactions with the chromatin. The blot was probed with anti-GCNA (GCNA-1), anti-H3 and anti-tubulin antibodies. Data is representative from three independent experiments. B. Schematic representation of constructs used in (C), (D), (E) and (F). C. Western blot of the FLAG immunoprecipitation. 3T3 cells were transiently transfected with FLAG-mGCNA and a FLAG immunoprecipitation was performed on the soluble fraction. The blot was probed with anti-FLAG, anti-H3 and anti-beta-actin antibodies. Data is representative from three independent experiments. D. Western blot of the FLAG immunoprecipitation. 3T3 cells were transiently transfected with FLAG-mGCNA and a FLAG immunoprecipitation was performed on the soluble fraction. The blot was probed with anti-FLAG, anti-H2B and anti-beta-actin antibodies. Data is representative from three independent experiments. E. Western blot of the GFP immunoprecipitation. 293-T cells were transiently transfected with GFP-fused proteins and a GFP immunoprecipitation was performed on the soluble fraction. The blot was probed with anti-GFP, anti-H3 and anti-beta-actin antibodies. Data is representative from three independent experiments. F. Western blot of the GFP immunoprecipitation. 293-T cells were transiently transfected with GFP-fused proteins and a GFP immunoprecipitation was performed on the soluble fraction. The blot was probed with anti-GFP, anti-H2B and anti-beta-actin antibodies. Data is representative from three independent experiments. G. Proximity ligation assay performed with anti-H3 and anti-GCNA (Tra98) antibodies on wild-type and *Gcna*^−^/Y adult testis sections. Data is representative from two independent experiments.

Histones are a major component of chromatin and their interaction with DNA relies on charged-based interactions. Since histones are highly basic and mGCNA is both highly acidic and associates with chromatin through electrostatic interactions, we hypothesised that mGCNA may interact with chromatin by binding to histones. We therefore tested if mGCNA physically interacts with histones by immunoprecipitation. As indirect chromatin interactions may confound our results, these experiments were conducted using the soluble, chromatin-free fraction. Additionally, samples were treated with nucleases. We observed that mGCNA can bind to H3 and H2B suggesting that mGCNA can bind the core histones H3-H4 and H2A-H2B (Fig 6B, C and D). We could not obverse any interaction between H1 and mGCNA, suggesting that the interaction between mGCNA and histones is specific to the core nucleosome components (Appendix Fig S7A). We then sought to test if human GCNA (hGCNA) also interacts with histones and found that it also interacts with H3 and H2B (Fig 6B, E and F). Moreover, we observed that the IDR of hGCNA, which is conserved with mGCNA, is sufficient for this interaction.

Next, we analysed if the interaction between mGCNA and core histones also occurred *in vivo*. In mice testes, we observed that GCNA co-localises with H3 in germ cells (Appendix Fig S7B). Furthermore, we observed by PLA that mGCNA is in close proximity to H3, from spermatogonia to round spermatids (Fig 6G). This result suggests that the chromatin association and histone binding of mGCNA occur in the germ line too.

### Mouse GCNA is a histone chaperone

Histones-binding proteins such as histone chaperones are characterised by highly acidic and disordered regions akin to that observed in mGCNA (Warren and Shechter, 2017). Given that mGCNA and hGCNA bind core histones, we sought to test if mGCNA exhibited histone chaperone activity. For this purpose, we purified N-terminally MBP-tagged mGCNA from human cells (Appendix Fig S8). A key property of histone chaperones is the ability to directly bind core histones. We therefore tested whether mGCNA could directly bind core histones. Limited proteolysis experiments showed increased protection of H3-H4 and H2A-H2B in presence of MBP-mGCNA compared to the MBP control (Fig 7A and B, lanes 4 and 8). These results suggest that mGCNA can directly interact with core histones and support the immunoprecipitation experiments presented earlier.

**Figure 7 -.**
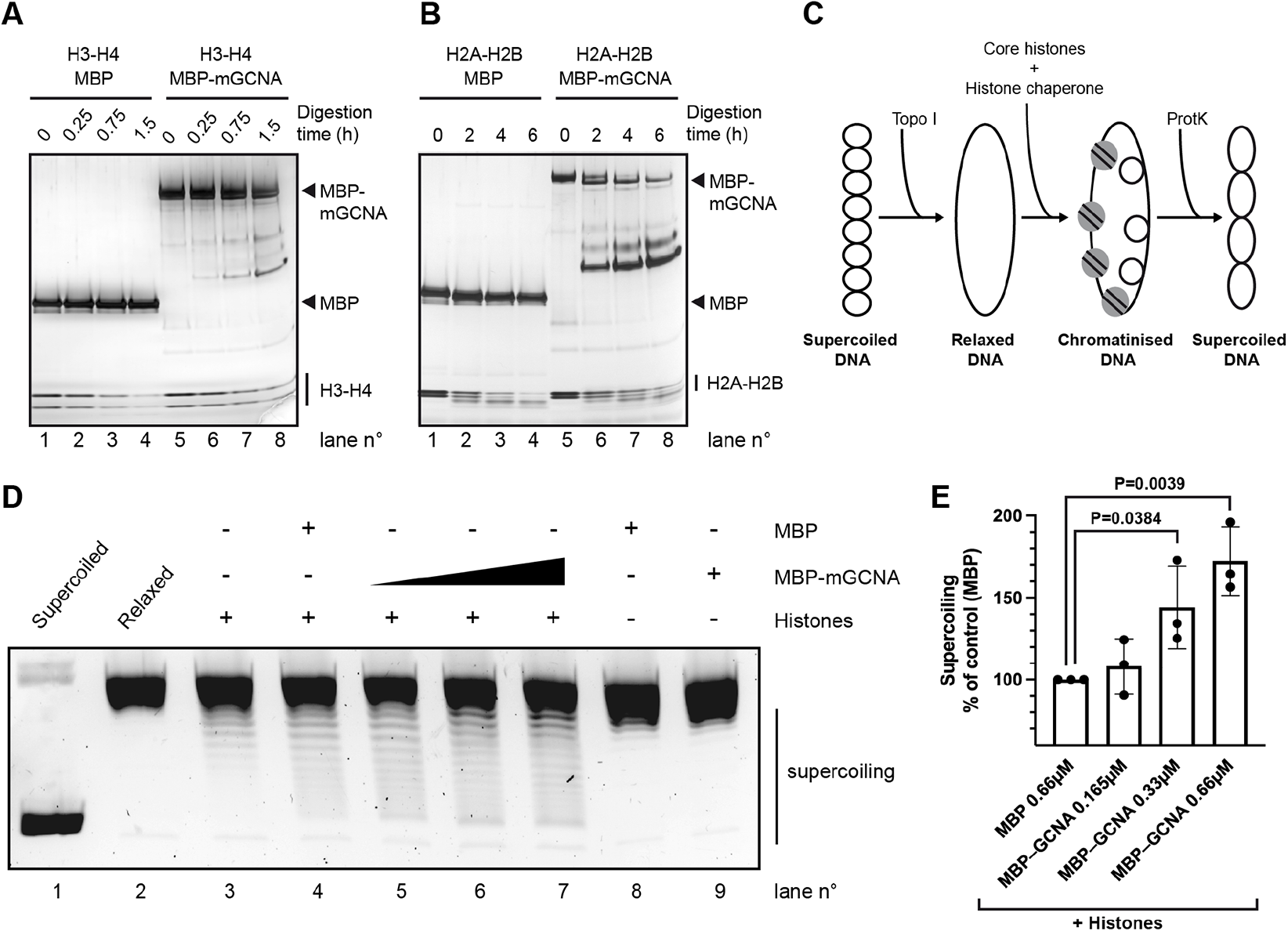
*In vitro* activities of mouse GCNA. A. Representative limited proteolysis assay performed with H3-H4, MBP and MBP-mGCNA. Proteins were incubated with trypsin for 0, 15, 45 and 90 min then analysed by SDS-PAGE followed by silver staining. Data is representative from three independent experiments. B. Representative limited proteolysis assay performed with H2A-H2B, MBP and MBP-mGCNA. Proteins were incubated with trypsin for 0, 2, 4 and 6 hours then analysed by SDS-PAGE followed by silver staining. Data is representative from two independent experiments. C. Schematic representation of the *in vitro* nucleosome assembly activity assay. A supercoiled plasmid is relaxed by TopoI then DNA was incubated with core histones and the histone chaperone. While nucleosomes are formed, supecoils are also formed into the plasmid and supercoiling is analysed by gel electrophoresis after deproteinisation of the plasmid. D. Representative *in vitro* nucleosome assembly activity assay with MBP and MBP-mGCNA. In this assay, MBP was used at a concentration of 0.66µM and MBP-mGCNA was used at 0.165, 0.33 or 0.66µM. Data is representative from three independent experiments. E. Quantification of the signal of the supercoiled plasmid in presence of histones and increasing concentrations of MBP-mGCNA (lanes 5 to 7 in (D)), relative to the signal with histones and MBP (lane 4 in (D)). Data represent the mean and S.D.. Data represent three independent experiments. *P* values were calculated by using an unpaired t-test.

We then sought to assess the potential of mGCNA to act as a histone chaperone. Using purified MBP-mGCNA, we performed an *in vitro* nucleosome assembly activity assay using MBP as a negative control (Fig 7C and D). In this assay, nucleosome assembly is monitored by the supercoiling of a plasmid DNA. MBP or MBP-mGCNA alone showed no supercoiling activity on the relaxed circular plasmid DNA (Fig 7D, lanes 8 and 9). Histones alone or with MBP showed little supercoiling activity on the plasmid DNA (Fig 7D, lanes 3 and 4). Interestingly, we observed that addition of increasing concentration of MBP-mGCNA to core histones progressively increased plasmid supercoiling (Fig 7D, lanes 5 to 7 and Fig 7E). The plasmid supercoiling is enhanced in presence of mGCNA, compared to the tag-only control, suggesting a specific activity of mGCNA (Fig 7D, lanes 4 and 7 and Fig 7E). These results indicate that mGCNA is able to promote core histone deposition onto DNA as a *bona fide* histone chaperone.

## Discussion

The expression of GCNA is restricted to germ line and embryonic stem cells in metazoans suggesting a role during germ cell and embryonic development. The presence of a SprT domain of GCNA, shared with the SPRTN protease, implicated GCNA in the repair of DNA-protein crosslinks (DPCs) to maintain genome stability. Indeed, studies in *C. elegans*, *Drosophila* and zebrafish provide evidence for a role of GCNA in the repair of DPCs in the germline and during early development (Bhargava et al., 2020; Dokshin et al., 2020). However, the absence of the SprT protease domain in mouse GCNA (mGCNA) suggests a function independent of DPC repair. Consistent with this, we have found that GCNA-deficient mESCs are not hypersensitive to DPC-inducing agents.

Although lacking the SprT domain, mGCNA shares a conserved intrinsically disordered region (IDR) with other GCNAs, including human GCNA. We therefore used the mouse protein as a model to explore these protease independent roles of GCNA. Our results show that mGCNA associates with chromatin, even in the absence of DNA damage. In addition, GCNA is highly acidic and predicted to be disordered akin to histone chaperones and we find that mGCNA directly binds core histones and also possesses histone chaperone activity *in vitro*. Moreover, we find that mGCNA is present at sites of nascent DNA synthesis, similar to other histone chaperones (Sirbu et al., 2011). Interestingly, GCNA interacts with components of the MCM2-7 complex in *Drosophila* suggesting that this association of GCNA with DNA replication is evolutionarily conserved (Bhargava et al., 2020). Consistent with DNA replication association and histone binding activity, we find that mGCNA ectopic expression causes alterations in the progression of DNA replication. Together, these data suggest that mGCNA is involved in chaperoning of histones during DNA replication.

Supporting this, we find that *in vivo*, the loss of GCNA leads to the accumulation of SSCs in S-phase. The accumulation of GCNA-deficient SSCs in this stage of the cell cycle could be explained by the loss of histone chaperone activity. Firstly, mGCNA may be important in chaperoning parental histones evicted by the MCM 2-7 complex and/or chaperoning newly synthesised histones, promoting their loading after the replication machinery (Fig 8, see (1)). Whether mGCNA acts to recycle histones or in *de novo* histone deposition remains to be determined. In the absence of mGCNA, parental and/or new histones may be less efficiently chaperoned, slowing fork progression or leading to aberrant positioning of histones, e.g. leading to regions of single-stranded DNA (Fig 8, see (1’)). Interestingly, in a recent study in *S.cerevisiae*, authors proposed that the SPRTN/GCNA homolog Wss1 targets the degradation of histones bound non-covalently to single-stranded DNA, preventing impaired fork progression (Maddi et al., 2020). Therefore, GCNA in rodents may have specialised in one ancestral function of the Wss1/SPRTN family which is the support of the management of histones during DNA replication.

**Figure 8 -.**
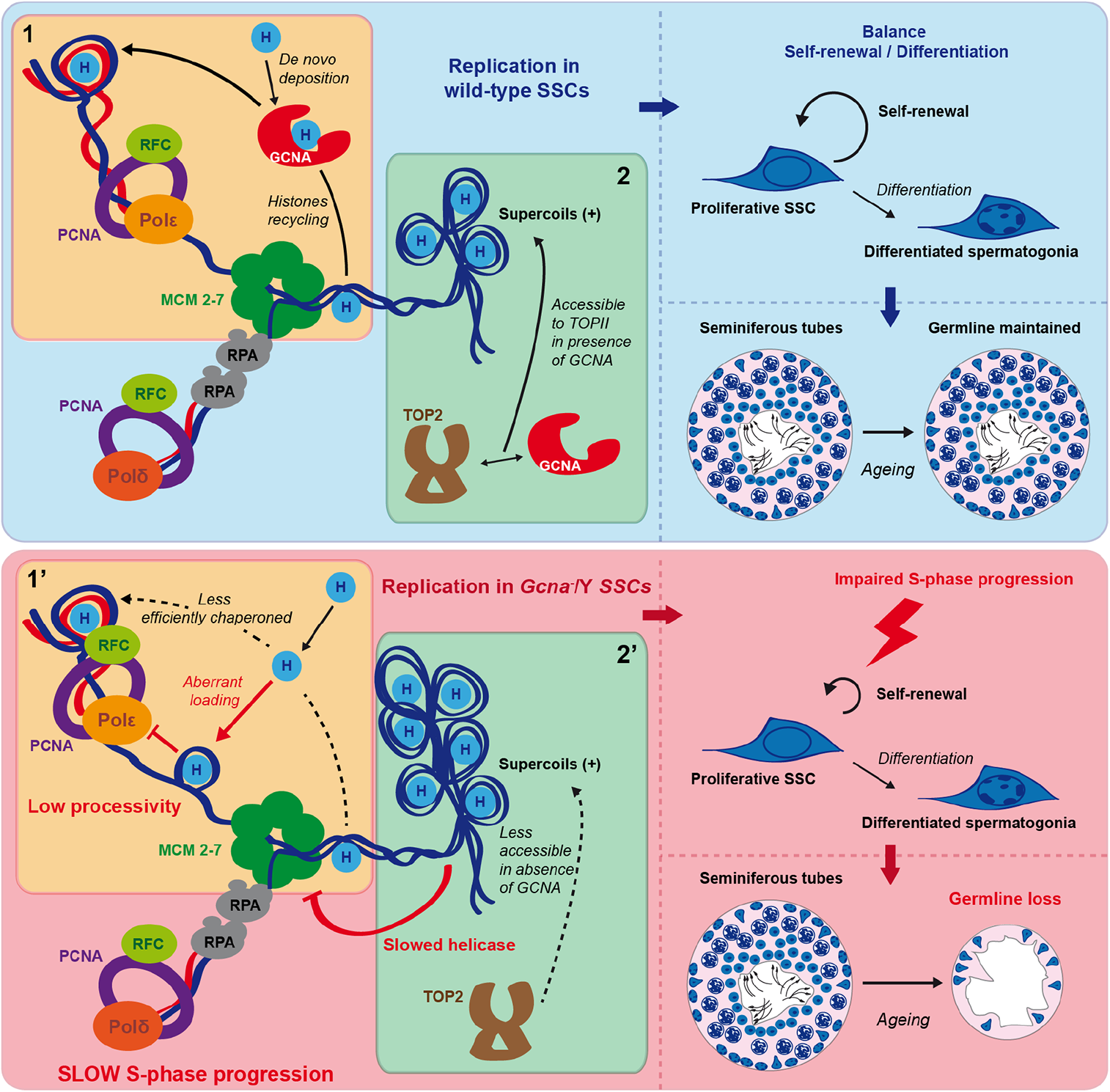
Working model of the putative roles of GCNA during murine spermatogenesis. In this model, two molecular roles of mGCNA are proposed (noted (1) and (2)). First, mGCNA is chaperoning histones that are then loaded after DNA synthesis (role (1)). Note that for simplification reasons, mGCNA is depicted as acting on the leading strand. Mouse GCNA could act as an histone chaperone on both strands, during DNA replication. Second, mGCNA interacts with TOP2 and supports relaxation of DNA supercoils ahead of the replication fork, in nucleosomes-rich regions (role (2)). Mouse GCNA supports the progression of S-phase in SSCs and therefore, supports their self-renewal. Consequently, SSCs are able to maintain homeostasis in the testis and mature sperm cells are produced through the mice life. However, in the absence of mGCNA, S-phase progression is slowed down because of a less efficient chaperoning of histones (see (1’)) or/and an accumulation of supercoils ahead of the replication fork (see (2’)). Thus, self-renewal of SSCs is impaired, leading to their loss. Consequently, SSC loss in the absence of GCNA triggers an age dependent reduction of the sperm production in males.

Alternatively, mGCNA may promote DNA replication through the interaction with TOP2. We observed that whilst mGCNA is dispensable for the repair of TOP2 crosslinked to DNA, GCNA does interact with TOP2. During replication, positive supercoils are formed ahead of the replication fork as the MCM 2-7 complex progresses. To prevent arrest of the replication fork due to high topological constraints, TOP2 relaxes the DNA (Ahmed and Dröge, 2020). However, in less accessible regions of chromatin, TOP2 may cooperate with a histone chaperone to have access to the DNA and we propose that GCNA could potentially help TOP2 at this step (Fig 8, see (2)). In the absence of mGCNA, TOP2 may have less access to DNA in nucleosome-rich regions, causing the replication fork to progress more slowly and impairing S-phase progression (Fig 8, see (2’)).

The loss of GCNA impacts male but not female germ cell production. GCNA-deficient male mice initially have intact spermatogenesis. However, this becomes compromised upon ageing. We hypothesised that this long-term failure to sustain spermatogenesis may be due to a defect in the SSC pool. This is consistent with a lack of defect during embryonic development. As adult females do not have an equivalent to SSCs, this may explain why the phenotypes are restricted to males. GCNA-deficient males initially have normal number of SSCs but this number decreases upon ageing, unlike in wild-type mice. This could be explained by either apoptosis, increased SSC differentiation, failure of SSC self-renewal, or a combination of these possibilities. We did not detect any increase in apoptosis or potentiated differentiation in the SSC pool of GCNA-deficient mice. However, in the absence of GCNA, aged SSCs show impaired S-phase progression. In mammals, the fine-tuned balance between self-renewal and differentiation of the SSCs allows the continuous production of gametes throughout life (Fig 8). As old GCNA-deficient SSCs replicate less due to impaired S-phase progression, they are more likely to be lost through normal differentiation rate. Consequently, the SSC pool is progressively depleted and the production of gametes is reduced in a time-dependent manner.

Murine GCNA represents an attractive separation-of-function model to interrogate its human homolog. Indeed, we observe that the IDR of human GCNA (hGCNA) shows similarities with its murine homolog, with a highly acidic region. Akin to the murine protein, hGCNA interacts with core histones. Furthermore, we observe that the IDR of hGCNA is sufficient for this interaction. However, hGCNA retains a SprT domain, like the invertebrate homologs. So, one could ask what is the molecular role of hGCNA in the germ line. We propose that hGCNA targets histone proteolysis thanks to the coupling of the binding activity of its IDR and the protease activity of the SprT domain.

Functionally, GCNA appears to be of crucial importance for the maintenance of the human male germ line. Indeed, two recent reports describe the phenotype of several azoospermic men carrying mutations in the *GCNA* locus (Arafat et al., 2021; Hardy et al., 2021). These mutations may lead to loss of GCNA protein but it is worth nothing that some mutations occur within the IDR. Strikingly, histological analysis of testis biopsies from two patients with *GCNA* mutations revealed a Sertoli-cell only phenotype in these two men (Hardy et al., 2021). This result strongly suggests that GCNA in humans may preserve the SSC pool, like we characterise here in mice. Therefore, we show here for the first time that the mouse model is relevant for the study of GCNA as it phenocopies mutations of GCNA in humans, despite the absence of the SprT domain in the mouse protein. Moreover, this result strongly suggests that the IDR of GCNA plays a critical role in the function of GCNA during mammalian spermatogenesis. Finally, we propose that GCNA in mammals is a histone binding protein required for SSC maintenance, through its roles in the support of DNA replication.

## Materials and methods

### Mice

All animal experiments undertaken in this study were approved by the Medical Research Council’s Laboratory of Molecular Biology animal welfare and ethical review body and the UK Home Office under the Animal (Scientific Procedures) Act 1986 (license no. PP6752216). All mice were maintained under specific pathogen-free conditions in individually ventilated cages (GM500; Techniplast) on Lignocel FS-14 spruce bedding (IPS) with environmental enrichment (fun tunnel, chew stick and Enviro-Dri nesting material (LBS)) at 19–23 °C with light from 7:00 a.m. to 7:00 p.m. and fed Dietex CRM pellets (Special Diet Services) ad libitum. No animals were wild and no field-collected samples were used. Unless otherwise stated, mice were maintained on a C57BL/6J background. *GOF18-GFP (Tg(Pou5f1-EGFP)2Mnn)* (MGI ID: 3057158) JAX (stock ID: 004654) mice were purchased from The Jackson Laboratory (Szabo et al., 2002). *Gcna*^tm1.1Dcp^ (*Gcna 2lox (2L))* (Carmell et al., 2016)(MGI ID: 5910931) JAX (stock ID: 031055) mice were purchased from The Jackson Laboratory. To generate the *Gcna* Δexon 4 allele, or *Gcna* knock-out allele, *Gcna*^tm1.1Dcp^ mice were bred with mice carrying the *Stella-Cre* allele, or *Dppa3*^tm1(cre)Peli^ (Liu et al., 2011)(MGI ID: 5004882). Genotyping of the *Gcna* knock-out allele was performed by PCR using the following primers: *Gcna* Fw (5’ GGATAGCAAAGGTTTATCAAC 3’) and *Gcna* Rv (5’ TGTGGTCCATAGCAAAATAAGG 3’) (Appendix Fig S1A and B). Gender of mice was determined by PCR as previously described (Clapcote and Roder, 2005).

### Assessment of the fertility of mice

To determine if GCNA-deficient mice were capable of giving rise to offspring, test mice were paired with wild-type C57BL/6J mates of the opposite sex. Females were examined daily for the presence of copulation plugs. The numbers of pups born over three successive months were recorded. Individuals performing the copulation plug checks were blinded to the genotype of mice.

### Epididymal sperm counts

One epididymis (caput, corpus and cauda) per mouse is placed in a 35 mm dish with 1 ml of PBS, pre-warmed at 37°C. Then, epididymes were punctuated with 19G needles. Dishes were then incubated at 37°C to let the spermatozoa swim-out. After 30 min, epididymes were flushed using the supernatant. Supernatants were collected and placed in clean, new, 2 ml tubes. Then, epididymes were flushed again with 400 µl of fresh PBS and supernatants were collected and added to the previous 2 ml tubes with supernatants. Collected spermatozoa, in approximatively 1.4 ml of PBS, were then counted using a Haemocytometer.

### Plasmids

Before use, all plasmids were validated by Sanger sequencing. To transiently express FLAG-mGCNA in mammalian cells and in cell-free systems, pcDNA3.1(+)-FLAG-mGCNA was generated by cloning FLAG-mGCNA cDNA into pcDNA3.1(+) (catalog no. V790-20, ThermoFisher Scientific) between KpnI and EcoRI sites. To express FLAG-mSPRTN in cell-free systems, pcDNA3.1(+)-FLAG-mSPRTN was generated by cloning FLAG-mSPRTN cDNA into pcDNA3.1(+) between KpnI and EcoRI sites. The plasmid pcDNA3.1(+)-FLAG-mSPRTN E113Q was generated by site directed mutagenesis from pcDNA3.1(+)-FLAG-mSPRTN. To transiently express EGFP-hGCNA or the EGFP-IDR of hGCNA in mammalian cells, the cDNAs of full length human GCNA or IDR hGCNA were cloned into pEGFP-C1 (Clontech) between BglII and SalI sites, to respectively make pEGFP-C1-hGCNA or pEGFP-C1-IDR hGCNA. The plasmid pEGFP-C1-hGCNA E593Q was generated by site directed mutagenesis from pEGFP-C1-hGCNA. To express mGCNA into *E.coli*, mGCNA cDNA was cloned into pOPT, between NdeI and BclI sites. In order to N-terminally tag mGCNA with MBP, mGCNA was first cloned into pOPTM between NdeI and BclI sites. Then, to transiently express MBP-mGCNA in Expi293 cells, MBP-mGCNA was cloned from pOPTM-mGCNA into pcDNA3.1(+) (catalog no. V790-20, ThermoFisher Scientific), between KpnI and EcoRI sites. To constitutively express FLAG-2xNLS-EGFP or FLAG-mGCNA in 3T3 cells, pExpress-LoxBsr-FLAG-2xNLS-EGFP and pExpress-LoxBsr-FLAG-mGCNA plasmids were made in two steps. First, cDNAs coding FLAG-2xNLS-EGFP and FLAG-mGCNA were cloned into pExpress between HindIII and EcoRV sites (Arakawa et al., 2001), to create pExpress-FLAG-2xNLS-EGFP and pExpress-FLAG-mGCNA. Those plasmids were after digested with ScaI and SpeI to separate the expression cassettes from the backbone. Then, the expression cassettes were cloned into pLoxBsr (Arakawa et al., 2001), between ScaI and SpeI, to create pExpress-LoxBsr-FLAG-2xNLS-EGFP and pExpress-LoxBsr-FLAG-mGCNA.

### Cell culture

Cells were cultivated in a 5% CO_2_ humidified incubator at 37°C. Mouse NIH3T3 cells (catalog no. CRL-1658, ATCC) and human HEK-293T cells (catalog no. CRL-3216, ATCC) were grown in DMEM (catalog no. 31966-021, ThermoFisher Scientific) supplemented with 10% Fetal Bovine Serum (catalog no. 10270-106, ThermoFisher Scientific). Expi293 cells (catalog no. A14527, ThermoFisher Scientific) were maintained in an 8% CO_2_ humidified incubator at 37°C, 125 RPM, and grown in Expi293 Expression medium (catalog no. A1435101, ThermoFisher Scientific). BAC9 mouse ESCs were a kind gift from A. Surani. Pluripotent mESCs were maintained in N2B27 media supplemented with the glycogen synthase kinase 3 inhibitor, CHIR 99021 (catalog no. 1386; Axon Medchem), the MAPK/ERK pathway inhibitor PD 0325901 (catalog no. 1408; Axon Medchem) and mouse Leukemia Inhibitory Factor (Cambridge Stem Cell Institute) referred to as to 2i + LIF media (Hayashi and Saitou, 2013). Mouse ESCs were maintained without feeders, on fibronectin-coated plates (catalog no. FC010, Sigma-Aldrich). Each cell line was regularly screened for the absence of mycoplasma contamination. Transfections were performed by using the polyethylenimine (PEI) method.

### Generation of 3T3 clones stably expressing FLAG-2xNLS-EGFP or FLAG-mGCNA

Mouse NIH3T3 cells were transfected with pExpress-LoxBsr-FLAG-2xNLS-EGFP or pExpress-LoxBsr-FLAG-mGCNA then later selected for resistance to Blasticidin (5 µg/ml, catalog no. A11139-03, Gibco). Resistant clones were then screened for the expression of FLAG-2xNLS-EGFP or FLAG-mGCNA by immunoblotting of FLAG (see later in “Western Blotting” part). For each constructs, two independent and successfully transfected clones were then expanded and maintained in culture with Blasticidin. To determine the growth rate, cells were counted at the indicated times using a cell counter (Vi-CELL XR Cell Viability Analyser, Beckman Coulter).

### Derivation of mESCs

*Gcna^+/-^* females were mated with wild-type males and the next day, embryos were collected and incubated in Embryomax KSOM Mouse embryo Media (catalog no. MR-020P-5F, Sigma-Aldrich) in a 5% CO_2_, 1% O_2_, humidified incubator at 37°C. After several days of culture, embryos hatched and were then placed on irradiated feeder mouse embryonic fibroblasts. Growing colonies of mESCs were then picked and mESC were expanded in 2i + LIF media (see previously, in “Cell culture” part), without feeders, on fibronectin-coated plates and in a 5% CO_2_ humidified incubator at 37°C. Clones were screened for their gender and genotype, as described earlier (see “Mice” part).

### Sensitivity to DPC-inducing agents

Sensitivity to DPC-inducing agents was determined by seeding 1000 mESCs per well of a 96-well flat-bottom plate, in supplemented KO-DMEM (KnockOut DMEM (catalog no. 10829-018, ThermoFisher Scientific), 12.5% Fetal Bovine Serum (catalog no. 10270-106, ThermoFisher Scientific), 0.1 mM β-mercaptoethanol (catalog no. 31350-010, ThermoFisher Scientific), 1X penicillin-streptomycin (catalog no. 15070-063, ThermoFisher Scientific), 1 mM Sodium Pyruvate (catalog no. 11360-070, ThermoFisher Scientific), 1X GlutaMAX supplement (catalog no. 35050-061, ThermoFisher Scientific), 1X Non Essential Amino Acids (catalog no. 11140-035, ThermoFisher Scientific)) with 10 ng/ml LIF(Cambridge Stem Cell Institute). The next day, cells were exposed to formaldehyde (catalog no. 28906, Thermo Scientific) or etoposide (catalog no. 2200, Cell Signaling Technology). Three days of culture post-exposure, the MTS cell viability reagent (CellTiter 96 AQueous One Solution Cell Proliferation Assay, catalog no. G3582, Promega) was added, plates were incubated at 37°C for 3 h and the absorbance at 490 nm was measured.

### Phylogenetic analysis of IDRs of metazoan GCNA proteins

IDRs of metazoans GCNA proteins were determined using GlobPlot 2 (Linding et al., 2003). Electrostatic charges of residues of these IDR were then determined using the EMBOSS charge software (https://www.bioinformatics.nl/cgi-bin/emboss/charge). Finally, representations of charges distributions in IDRs were obtained using the Heatmapper software (Babicki et al., 2016).

### Protein purification

#### Purification of MBP-mGCNA

Mouse GCNA N-terminally fused to MBP was purified from Expi293 cells as follow: 3 _x_10^9^ Expi293 cells were transfected with pcDNA3.1(+)-MBP-mGCNA by PEI transfection (Fang et al., 2017). Cells were then incubated in presence of 3.5 mM valproic acid (catalog no. P4543, Sigma-Aldrich) in culture media for 4 days at 37°C. Cells were then pelleted at 300g for 5 min and lysed in lysis buffer (20 mM Tris ph 7.4, 200 mM NaCl, 1 mM EDTA, 1X Protease inhibitors (catalog no. 11873580001, Roche), 0.5% Triton X-100, 5 mM CaCl_2_, 20 µg/ml RNAse A (catalog no. R6148, Sigma-Aldrich), 1000 U/ml MNase (catalog no. M0247S, New England Biolabs)) at a ratio of 1 ml of lysis buffer per 100 mg of cell pellet. Cells were sheared with a 21G needle then sonicated for 10 min at 4°C with a non-contact sonicator (30 sec ON/30 sec OFF cycles, 40% amplitude, catalog no. Vibra-Cell VC 750, Sonics & Materials). Extracts were then spun at 16,200g for 20 min at 4°C and supernatants were collected. Cleared extracts were diluted ten times in column buffer (20 mM Tris ph 7.4, 200 mM NaCl, 1 mM EDTA, 1X Protease inhibitors (catalog no. 11873580001, Roche)). 1 L of diluted extracts were incubated with 9 ml of packed amylose resin (catalog no. E8021S, New England Biolabs) and incubated at 4°C for 2h with gentle rotation. Extracts with resin were then loaded onto a column and washed with 3 volumes of column buffer, then 6 volumes of column buffer with 1M NaCl and with 3 volumes of elution buffer without Maltose (25 mM Tris ph 7.4, 10% glycerol, 75 mM NaCl, 2 mM DTT, 1 mM EDTA). Proteins were then eluted with 25 ml of elution buffer supplemented with 10 mM Maltose and several fractions were made. Fractions were then analysed by SDS-PAGE and fractions containing MBP-mGCNA were pooled. Pooled fractions were loaded onto an Anion exchange Maxi column (catalog no. 78243, ThermoScientific). The column was washed three times with buffer A (25 mM Tris pH 7.4, 75 mM NaCl, 10% glycerol, 1 mM EDTA, 2 mM DTT), 10 ml per wash. Then, the column was serially washed with buffer A containing increasing concentrations of NaCl (100, 150, 200, 300 mM NaCl), 10 ml per wash. Then, MBP-mGCNA was eluted three times with 2 ml buffer A containing 500 mM NaCl. Elutions with pure MBP-mGCNA were aliquoted and stored at −80°C.

#### Purification of MBP

MBP was purified from *E.coli* as follow: Rosetta (DE3) cells (catalog no. 70954, Sigma-Aldrich) transformed with pOPTM were incubated ON at 37°C in SOB media (2% Tryptone, 0.5% Yeast extract, 10 mM NaCl, 2.5 mM KCl, 10 mM MgCl_2_) containing chloramphenicol (25 µg/ml) and ampicillin (100µg/ml). The next day, 5 ml of ON culture was diluted in 500 ml 2X TY media (16 g/L Tryptone, 10 g/L yeast extract, 5 g/L NaCl) containing chloramphenicol and ampicillin and the culture was incubated for 4h at 37°C. Then, 500 ml 2X TY media with chloramphenicol and ampicillin and 1 mM IPTG were added to the culture. The culture was then incubated for 15h at 20°C. Culture was then pelleted and stored at −80°C. The cell pellet was then lysed with B-PER buffer (catalog no. 78243, ThermoFisher Scientific) according to the manufacturer’s instructions. Lysate was then spun at 16,200g for 15 min at 4°C and supernatant was collected. The supernatant was then diluted ten times in column buffer (20 mM Tris ph 7.4, 200 mM NaCl, 1 mM EDTA, 1X Protease inhibitors (catalog no. 11873580001, Roche)). Diluted lysate was then incubated with 9 ml of packed amylose resin (catalog no. E8021S, New England Biolabs) and incubated at 4°C for 2h with gentle rotation. Extracts with resin were then loaded onto a column and washed with 3 volumes of column buffer, then 6 volumes of column buffer with 1M NaCl and with 3 volumes of elution buffer without Maltose (25 mM Tris ph 7.4, 10% glycerol, 75 mM NaCl, 2 mM DTT, 1 mM EDTA). Proteins were then eluted with 25 ml of elution buffer supplemented with 10 mM Maltose and several fractions were made. Fractions were then analysed by SDS-PAGE and fractions with MBP were pooled. The buffer containing MBP was then exchanged with the storage buffer of MBP-mGCNA (25 mM Tris pH 7.4, 500 mM NaCl, 10% glycerol, 1 mM EDTA, 2 mM DTT) using a Vivaspin-20 concentrator (catalog no. VS2021, Sartorius). Five buffer exchanges with the storage buffer were performed and purified MBP was then aliquoted and stored at −80°C.

### Limited proteolysis assay

In a 100 µl reaction, 1.4 µg of H3-H4 (catalog no. M2509S, New England Biolabs) or H2A-H2B (catalog no. M2508S, New England Biolabs) were incubated with 2.5 µg of MBP or MBP-mGCNA in Binding buffer (25 mM Tris pH 7.5, 150 mM NaCl, 5 % glycerol). Then, 0.5 ng of Trypsin Gold (for the proteolysis with H3-H4, catalog no. V528A, Promega) or 1 ng of Trypsin Gold (for the proteolysis with H2A-H2B) were added to the 100 µl reactions. Aliquots of 20 µg were then collected at the indicated time points and later analysed by SDS-PAGE. Polyacrylamide gels were then stained by using the Silver stain method (catalog no. 24612, ThermoFisher Scientific) and stained gel were imaged by using a ChemiDoc MP Imaging System, Biorad.

### Plasmid supercoiling assay

The circular plasmid, phiX174 RF1 (catalog no. SD0031, ThermoFisher Scientific) was pretreated for 24h at 37°C with Topoisomerase I (5U of Topoisomerase I per µg of plasmid, catalog no. 38042-024, Invitrogen) in 50 mM Tris pH 7.5, 50 mM KCl, 10 mM MgCl_2_, 0.5 mM DTT, 0.1 mM DTT, 0.1 mM EDTA, 30 µg/ml BSA. Histones H3-H4 (0.33 µM, catalog no. M2509S, New England Biolabs) and H2A-H2B (0.33 µM, catalog no. M2508S, New England Biolabs) were incubated with increasing concentrations of MBP-mGCNA (0.165, 0.33 or 0.66 µM) or with MBP (0.66 µM) in 15 µl reaction containing 20 mM Tris ph 7.4, 0.5 mM EDTA, 10% glycerol, 0.1 mg/ml BSA, 0.5 mM DTT. NaCl in the reaction was provided by the protein storage buffers added into the reaction with the proteins and the concentration was set at 216 mM for each reaction. Proteins were then incubated at 37°C for 30 min. Then, 100 ng of relaxed phiX174 was added into each chaperone-histones mix and incubated for 1h at 37°C. Reactions were then stopped by adding 16 µl of stop buffer (25% glycerol, 60mM Tris pH 8, 30 mM EDTA, 2% SDS, 2 mg/mL proteinase K) and incubated for 1h at 37°C. Products were then analysed by 1% agarose gel electrophoresis in TBE buffer followed by SYBR Gold staining (catalog no. S11494, ThermoFisher Scientific). Gels were imaged with a ChemiDoc MP Imaging System, Biorad. Supercoilling was quantified using Fiji software (Schindelin et al., 2012). Each lane were manually selected with the software (Gels > Select First Lane, then Gels > Select Next Lane) and signal intensity for each lane was quantified by Fiji (Gels > Plot Lanes, then use the Wand tool to obtain the quantifications).

### Protein extracts preparation for Western Blotting

Mouse ESC were lysed in ice cold RIPA buffer (25 mM Tris-Hcl pH 7.4, 150 mM NaCl, 0.1% SDS, 0.5% Sodium Deoxycholate, 1% Triton X-100) complemented with protease inhibitors cocktail (catalog no. 11873580001, Roche) and let chill on ice for 10 min. Then, extracts were sonicated for 10 min at 4°C with a non-contact sonicator (30 sec ON/30 sec OFF cycles, 40% amplitude, catalog no. Vibra-Cell VC 750, Sonics & Materials). Extracts were spun at 16,200g for 10 min at 4°C, supernatants were collected and analysed by Western Blot. Testes were processed in the same way but were first homogenised in RIPA buffer with protease inhibitors using a Dounce Homogeniser (10 strokes with loose pestle then 10 strokes with the tight one).

### Immunoprecipitation

Cells were lysed in CSK buffer (10 mM PIPES pH 7.5, 100 mM NaCl, 300 mM Sucrose, 3 mM MgCl_2_) with 0.5 % Triton X-100 and protease inhibitors cocktail (catalog no. 11873580001, Roche) and let chill on ice for 10 min. Cells were then spun at 845g for 5 min at 4°C and the supernatant containing the cellular soluble fraction was collected. One volume of soluble fraction was then diluted with 5 volumes of Binding buffer (20 mM Tris-Hcl pH 7.4, 150 mM NaCl, 10% glycerol, 1 mM CaCl_2_, 1 mM MgCl_2_, 0.1% Bovine serum albumin, 1 mM β-mercaptoethanol, protease inhibitors cocktail (catalog no. 11873580001, Roche)) then 90 U of Benzonase (catalog no. 70664, Merk Millipore) and 3 U of DNAse I (catalog no. M0303, New England BioLabs) were added per ml of mix. For each immunoprecipitation, 25 µl of anti-FLAG (catalog no. M8823, Sigma-Aldrich) or anti-GFP (catalog no. gtma-20, Chromotek) magnetic beads were then washed with PBS + 0.05% Tween 20 and then with Binding buffer. After washes, magnetic beads were added to the cellular soluble fraction diluted in Binding buffer. The immunoprecipitation mix was then incubated ON at 4°C with gentle rotation. The next day, magnetic beads were washed three times with CSK buffer + 0.5 % Triton X-100 and protease inhibitors cocktail, 5 min with gentle rotation at 4°C for each wash. Elution was then done by adding 1.5 X LDS buffer (catalog no. NP0007, ThemoFisher Scientific) containing 5 % β-mercaptoethanol to the beads. Samples were then analysed by Western Blot.

### Cellular fractionation for chromatin isolation

Cells were lysed in one volume of CSK buffer (10 mM PIPES pH 7.5, 100 mM NaCl, 300 mM Sucrose, 3 mM MgCl_2_) with 0.5 % Triton X-100 and protease inhibitors cocktail (catalog no. 11873580001, Roche) and let chill on ice for 10 min. Cells were then spun at 845g for 5 min at 4°C and the supernatant containing the cellular soluble fraction was collected and stored. The pellet containing the chromatin fraction was washed two times with three volumes of CSK buffer containing 0.5 % Triton X-100 and protease inhibitors cocktail. For DNAse I digestion of the chromatin fraction, chromatin was incubated in DNAse I buffer (10 mM Tris-Hcl pH 7.6, 2.5 mM MgCl_2_, 0.5 mM CaCl_2_) with 20 U of DNAse I for 45 min at 37°C. After digestion, the chromatin fraction was then washed two times with five volumes of CSK buffer containing 0.5 % Triton X-100 and protease inhibitors cocktail. The chromatin fraction was finally resuspended in 1.5 X LDS buffer (catalog no. NP0007, ThemoFisher Scientific) containing 5% β-mercaptoethanol and sonicated for 10 min at 4°C with a non-contact sonicator (30 sec ON/30 sec OFF cycles, 40% amplitude, catalog no. Vibra-Cell VC 750, Sonics & Materials). Samples were then analysed by Western Blot.

### Chromatin isolation on synchronised cells

NIH3T3 cells were synchronised in S-phase as follow: cells were plated at a density of 5 x10^5^ cells per 10 cm dish. Twenty-four hours after plating, cells were synchronised by being washed two times with DMEM (catalog no. 31966-021, ThermoFisher Scientific) and then placed in DMEM supplemented with 0.5% Fetal Bovine serum (catalog no. 10270-106, ThermoFisher Scientific). Cells were incubated for 48h and then released from serum starvation by incubating them for 15 h in DMEM supplemented with 10% Fetal Bovine serum. After 15 h of culture, cells in S-phase were collected and processed for chromatin isolation as described earlier.

### Chromatin isolation and NaCl gradient washes

Mouse embryonic stem cells were lysed in one volume of CSK buffer (10 mM PIPES pH 7.5, 100 mM NaCl, 300 mM Sucrose, 3 mM MgCl_2_) containing 0.5 % Triton X-100 and protease inhibitors cocktail (catalog no. 11873580001, Roche) and let chill on ice for 10 min. Cells were then spun at 845g for 5 min at 4°C and the supernatant containing the cellular soluble fraction was collected and stored. The pellet containing the chromatin fraction was washed two times with three volumes of CSK buffer (with modified NaCl concentration ranging from 100 mM to 700 mM) + 0.5 % Triton X-100 and protease inhibitors cocktail. Then, chromatin fraction was washed two times with the regular CSK buffer containing 0.5 % Triton X-100 and protease inhibitors cocktail. The chromatin fraction was finally resuspended in 1.5 X LDS buffer (catalog no. NP0007, Themo Fisher Scientific) containing 5% β-mercaptoethanol and sonicated for 10 min at 4°C with a non-contact sonicator (30 sec ON/30 sec OFF cycles, 40% amplitude, catalog no. Vibra-Cell VC 750, Sonics & Materials). Samples were then analysed by Western Blot.

### DNA pulldown assay

DNA pulldown assays were performed as follow. The HPLC-purified biotinylated 59 bp long oligonucleotide, 5’-GAT CTG CAC GAC GCA CAC CGG ACG TAT CTG CTA TCG CTC ATG TCA ACC GCT CAA GCT GC-3’-biotin-TEG, and its complementary oligonucleotide were used for the pulldown assay. Double strand hybridization was performed in 50 mM NaCl, 25 mM Tris-HCl, pH 7.5 buffer with the biotinylated oligonucleotide and its complementary oligonucleotide at 1:1.5 ratio by denaturing for 3 minutes at 95°C and allowing a slow progressive return to room temperature. Mouse recombinant FLAG-GCNA and FLAG-mSPRTN E113Q were produced in a cell-free system. The TnT T7 Quick Coupled Transcription/Translation System (catalog no. L1170, Promega) was used with pcDNA3.1(+)-FLAG-mGCNA and pcDNA3.1(+)-FLAG-mSPRTN E113Q plasmids in a 40 μl reaction according to the manufacturer’s instructions. Ten microliters of recombinant FLAG-mGCNA or FLAG-mSPRTN E113Q from cell-free extracts and 500 μg of Dynabeads M-280 Streptavidin (catalog no. 11205D, Themo Fisher Scientific) with immobilized ssDNA or dsDNA were incubated in 440 μl binding buffer (25 mM Tris-HCl, pH 7.5, 100 mM NaCl, 1 mM EDTA, 5 mg/ml BSA, 0.05% Tween 20, 10% glycerol, 1 mM β-mercaptoethanol, protease inhibitors cocktail (catalog no. 11873580001, Roche)). The DNA-protein mixture was incubated for 2 h at 4°C with gentle rotation. After incubation, magnetic beads were washed twice in 500 μl binding buffer without BSA and then washed once in 500 μl rinsing buffer (25 mM Tris-HCl, pH 7.5, 150 mM NaCl, 1 mM EDTA 0.05% Tween 20, 10% glycerol, 1 mM β-mercaptoethanol, protease inhibitors cocktail). Bound proteins were eluted with 1.5 X LDS (catalog no. NP0007, Themo Fisher Scientific) containing 5% β-mercaptoethanol. Samples were then analysed by Western Blot.

### IPOND

For each point, 4.5×10^6^ BAC9 mES cells were seeded in a 15 cm Petri Dish with 45 ml of supplemented KO-DMEM (KnockOut DMEM (catalog no. 10829-018, ThermoFisher Scientific), 12.5% Fetal Bovine Serum (catalog no. 10270-106, ThermoFisher Scientific), 0.1 mM β-mercaptoethanol (catalog no. 31350-010, ThermoFisher Scientific), 1X penicillin-streptomycin (catalog no. 15070-063, ThermoFisher Scientific), 1 mM Sodium Pyruvate (catalog no. 11360-070, ThermoFisher Scientific), 1X GlutaMAX supplement (catalog no. 35050-061, ThermoFisher Scientific), 1X Non Essential Amino Acids (catalog no. 11140-035, ThermoFisher Scientific)). After 48h of culture, IPOND was then performed as published (Sirbu et al., 2012).

### Western Blotting

Protein samples were supplemented with LDS buffer (catalog no. NP0007, Themo Fisher Scientific) and 5% β-mercaptoethanol final, boiled for 5 min at 95°C and resolved by polyacrylamide gel electrophoresis on NuPAGE 4 to 12%, Bis-Tris, Mini Protein gels (catalog no. NP0321BOX, ThermoFisher Scientific) in MOPS-SDS buffer (50 mM MOPS, 50 mM Tris base, 3.47 mM SDS, 1mM EDTA). Separated proteins were transferred onto 0.2 µm Nitrocellulose membranes (catalog no 10600015, GE Healthcare) in Tris-Glycine (25 mM Tris, 192 mM Glycine, ph 8.3) buffer with 20% ethanol. Transfer was set at 35 V for 1h30 in a Xcell II Blot module (catalog no. EI9051, ThermoFisher Scientific). After transfer, membranes were incubated for 1h in blocking buffer (Tris-buffered saline, 0.1% Tween 20, 5% non-fat dry milk). Then, membranes were incubated ON at 4°C in blocking buffer with the following antibodies: anti-GCNA clone GCNA-1 (1:1000, catalog no. 10D9G11, DSHB); anti-GCNA clone Tra98 (1:1000, catalog no. ab82527, Abcam); anti-histone H3 (1:7500, catalog no. ab1791, Abcam); anti-histone H2B (1:1000, catalog no. ab1790, Abcam); anti-α-Tubulin (1:3000, catalog no. T6199, Sigma-Aldrich); anti-FLAG (1:1000, catalog no. 2368, Cell Signaling Technology); anti-GFP (1:1000, catalog no. GF090R, Nacalai USA); anti-beta-Actin (1:2000, catalog no. ab8227, Abcam); anti-MBP (1:10,000, catalog no. E8032S, New England Biolabs); anti-CCNA2 (1:1000, catalog no. ab181591, Abcam); anti-VINCULIN (1:2000, catalog no. ab129002, Abcam); anti-H3 pSer10 (1:1000, catalog no. 9701S, Cell Signaling Technology); anti-PCNA (1:1000, clone PC10, catalog no. MABE288, Sigma-Aldrich); anti-PLZF (1:100, catalog no. sc-28319, Santa Cruz Biotechnology); anti-WT1 (1:1000, catalog no. ab89901, Abcam); anti-LAMINB1 (1:3000, catalog no. ab16048, Abcam); anti-Nanog (1:1000, catalog no. 8822, Cell Signaling Technology); anti-TOP2 (1:1000, catalog no. ab109524, Abcam); anti-SYCP3 (1:500, catalog no. ab15093, Abcam), anti-GroEL; anti-histone H1 (1:500, catalog no. ab61177, Abcam). Membranes were washed with TBS + 0.1% Tween 20 and then incubated for 1h at RT with the following secondary antibodies diluted in blocking buffer: swine anti-rabbit Ig HRP-conjugated (1:3000, catalog no. P0399, Dako) or goat anti-mouse Ig HRP-conjugated (1:5000, catalog no. P0447, Dako) or goat anti-rat IgG HRP-conjugated (1:2000, catalog no. 7077, Cell Signaling Technology). Membranes were then washed with TBS + 0.1% Tween 20. Then, membranes were incubated with an ECL Western Blotting Detection reagent (catalog no. RPN2106, GE Healthcare). Acquisition of the chemiluminescent signal was done using X-Ray films (catalog no. FM024, Photon Imaging Systems Ltd).

### Immunofluorescence on cultured cells

NIH3T3 cells were seeded on no. 1.5 coverslips (catalog no. 631-0150, VWR) 24h before experiments. If needed, cells were then exposed for 30 min with 10 µM EdU (catalog no. A10044, ThermoFisher Scientific). For the classic fixation, cells were washed twice for 5 min with PBS supplemented with 500 μM MgCl_2_ and 0.5 μM CaCl_2_ (PBS-S) then fixed with 4% paraformaldehyde (catalog no. 43368, Alfa Aesar) for 20 min and washed with PBS-S twice for 5 min. If cells were previously exposed to EdU, paraformaldehyde was replaced with formaldehyde (catalog no. 28906, Thermo Scientific). Cells were then permeabilised for 10 min with PBS-S containing 0.1% Triton X-100 and washed PBS-S twice for 5 min.

For the pre-extraction before fixation, cells were washed once with PBS-S for 3 min, then with CSK buffer (10 mM Pipes, pH 7.0, 100mM NaCl, 300 mM sucrose, 3 mM MgCl_2_) for 3 min and incubated twice for 3 min with CSK buffer containing 0.5% Triton X-100 and complete protease inhibitors cocktail (catalog no. 11873580001, Roche). Cells were after washed with PBS-S, fixed with 4% paraformaldehyde (catalog no. 43368, Alfa Aesar) for 20 min and washed with PBS-S. If cells were previously exposed to EdU, paraformaldehyde was replaced with formaldehyde (catalog no. 28906, Thermo Scientific).

Labelling of EdU was then performed using the Click-iT Plus EdU Cell Proliferation Kit, Alexa Fluor 594 dye (catalog no. C10639, ThermoFisher Scientific). After fixation and washes and EdU labelling, cells were blocked with PBS-S/0.1% Tween 20 (PBS-S-T) containing 5% BSA for 30 min. Cells were then incubated ON at 4°C with the following primary antibodies diluted in blocking buffer: anti-FLAG (1:200, M2 clone, catalog no. F1804, Sigma-Aldrich); anti-FLAG (1:200, catalog no. 2368, Cell Signaling Technology); anti-LAMIN B1 (1:500, catalog no. ab16048, Abcam); anti-RPA2 (1:400, catalog no. 2208, Cell Signaling Technology). Cells were then washed with PBS-S-T and incubated for 1h at 37°C with the following secondary antibodies diluted in blocking buffer: goat anti-rat Alexa Fluor 488 (1:500, catalog no. A-11006, ThermoFisher Scientific), goat anti-rabbit Alexa Fluor 594 (1:500, catalog no. A-11037, ThermoFisher Scientific), goat anti-mouse Alexa Fluor 488 (1:500, catalog no. A-11029, ThermoFisher Scientific). After washes with PBS-S-T, coverslips were incubated for 10 min with 2 μg/ml DAPI in PBS-S. After washes in PBS-S, coverslips were mounted on glass slides using Prolong Gold Antifade Mountant (catalog no. P36934, ThermoFisher Scientific). Images were captured with a LSM 780 confocal microscope (ZEISS).

### Histological analysis

Tissues were fixed in 10% neutral-buffered formalin (catalog no. HT5011-15ml, Sigma-Aldrich) or in Bouin’s solution (catalog no. 320700-1000, RAL Diagnostics) for 24-48 h and transferred in 70% ethanol. Fixed samples were embedded in paraffin and 3.5 μm sections cut, deparaffinised, rehydrated and stained with hematoxylin and eosin (H&E) following standard methods. Images were captured with an Axioplan 2 microscope (ZEISS).

### Immunofluorescence on testis sections

When the experiment involved the incorporation of EdU into germ cells, mice were given a single intraperitonal injection of EdU (catalog no. A10044, ThermoFisher Scientific), 50 mg/kg at 10 ml/kg. Four hours after injections, testes were biopsied, fixed in formalin (catalog no. HT5011-15ml, Sigma-Aldrich) and processed for histological analysis as described earlier. Sections of formalin-fixed, paraffin-embedded samples were deparaffinised and rehydrated following standard procedure. Slides were boiled for 20 min into antigen retrieval buffer (10 mM sodium citrate, pH 6.0). Slides were allowed to chill at room temperature then slides were washed three times in water for 5 min and then once for 5 min in Tris-buffered saline (TBS) containing 0.1% Tween 20. If needed, labelling of EdU was then performed using the Click-iT Plus EdU Cell Proliferation Kit, Alexa Fluor 594 dye (catalog no. C10639, ThermoFisher Scientific). Sections were then incubated in blocking buffer (TBS, 0.1% Tween 20, 5% Fetal Bovine Serum) for 1h at room temperature. Sections were incubated ON at 4°C with the following primary antibodies diluted in blocking buffer: anti-GCNA clone GCNA-1 (1:100, catalog no. 10D9G11, DSHB); anti-GCNA clone Tra98 (1:300, catalog no. ab82527, Abcam); anti-human GCNA (1:20, catalog no. HPA023476, Sigma-Aldrich); anti-LAMIN B1 (1:500, catalog no. ab16048, Abcam); anti-histone H3 (1:500, catalog no. ab1791, Abcam); anti-PCNA (1:400, catalog no. 13110, Cell Signaling Technology); anti-Cleaved Caspase 3 (1:300, catalog no. 9661, Cell Signaling Technology); anti-DNMT3B (1:250, catalog no. ab122932, Abcam); anti-KI67 (1:100, catalog no. M3062, Spring Bioscience); anti-CCNA2 (1:400, catalog no. ab181591, Abcam); anti-pH3, Ser10 (1:200, catalog no. 9701, Cell Signaling Technology); anti-PLZF (1:100, catalog no. sc-28319, Santa Cruz Biotechnology); anti-PLZF (1:300, catalog no. sc-22839, Santa Cruz Biotechnology), anti-γH2AX (1:200, catalog no. 2577, Cell Signaling Technology). Slides were washed three times for 5 min with TBS + 0.1% Tween 20 and then incubated for 1h at RT with the following secondary antibodies diluted in blocking buffer: goat anti-rat Alexa Fluor 488 (1:500, catalog no. A-11006, ThermoFisher Scientific), goat anti-rabbit Alexa Fluor 594 (1:500, catalog no. A-11037, ThermoFisher Scientific), goat anti-mouse Alexa Fluor 488 (1:500, catalog no. A-11029, ThermoFisher Scientific). After incubation with secondary antibodies, slides were washed three times in TBS + 0.1% Tween 20 for 5 min. After washes, slides were incubated for 10 min with 2 μg/ml DAPI in TBS then washed three times in TBS for 5 min. Slides were then briefly dipped into water and mounted with Prolong Gold Antifade Mountant (catalog no. P36934, ThermoFisher Scientific). Images were captured with a LSM 780 confocal microscope (ZEISS). Human testis sections (catalog no. 11-701 YC1, ProSci Inc.) were processed on the same way than mice testis sections.

### Proximity ligation assay (PLA) on testis sections

Deparaffinisation, rehydratation and antigen retrieval of formalin-fixed testis sections were done as described earlier. Then, PLA was proceed using the Duolink In Situ Red Starter Kit Mouse/Rabbit (catalog no. DUO92101, Sigma-Aldrich) following manufacturer’s instructions. Primary antibodies used in this study were anti-GCNA (1:300, Tra98 clone, catalog no. ab82527, Abcam), anti-histone H3 (1:500, catalog no. ab1791, Abcam) and anti-PCNA (1:400, catalog no. 13110, Cell Signaling Technology). The secondary antibody Duolink In Situ PLA Probe Anti-Mouse MINUS from the PLA kit was replaced with a Donkey anti-rat antibody (catalog no 712-005-150, Jackson ImmunoResearch) coupled to a MINUS probe using the Duolink In Situ Probemaker MINUS kit (catalog no. DUO92010, Sigma-Aldrich). Images were captured with a LSM 780 confocal microscope (ZEISS).

### Image analysis

Image were analysed using Fiji software (Schindelin et al., 2012). To quantify the amount of fluorescence per cell, cell nuclei were manually selected with the software and the fluorescence (IntDen) was quantified by Fiji (Analyse > Measure). Profiles of fluorescence intensity were obtained by first drawing a line on the region of interest then by quantifying the fluorescence intensity with Fiji (Analyse > Plot Profile).

### Quantification of Primordial Germ Cells

Urogenital ridges of E12.5 embryos carrying the GOF18-GFP reporter were isolated and placed into 150 μl of trypsin solution (2.5 μg/ml trypsin (Gibco), 25 mM Tris, 120 mM NaCl, 25 mM KCl, 25 mM KH2PO4, 25 mM glucose, 25 mM EDTA, pH 7.6) pre-warmed to 37 °C and incubated for 10 min at 37 °C. Subsequently, 1 μl of Benzonase endonuclease (catalog no. 70664, Merk Millipore) was added per sample and samples were disaggregated by gentle pipetting and incubated for a further 5 min at 37 °C. The trypsin was inactivated by adding 1 ml of PBS + 5% Fetal Bovine Serum (FBS). Following 10 min of centrifugation at 3,300 r.p.m., the cell pellet was resuspended in 100 μl of Alexa Fluor 647-conjugated anti-human/mouse SSEA1 antibody (catalog no. MC-480; BioLegend) diluted 1:100 in PBS + 2.5% FBS and incubated at room temperature for 10 min. Then, 300 μl of PBS + 2.5% FBS were added to the cell suspension and samples were immediately run on an ECLIPSE analyzer (Sony Biotechnology) and the data analysed using FlowJo v.10.1r5 (FlowJo LLC).

### Statistical analysis

The number of independent biological samples and technical repeats are indicated in the figure legends. Unless otherwise stated, data are shown as the mean ± S.D.. Analysis was performed in Prism 9 (GraphPad Software).

## Acknowledgements

We would like to thank Jessica McCool, Claire Knox, Liam Bray, Lucy Tredgett, Chloe Watson, the Ares and Biomed staff for assistance with animal procedures and experiments. We thank the Human Research Tissue Bank (National Institute for Health Research Cambridge Biomedical Research Centre) for processing the histology. We thank Azim Surani for the gift of the BAC9 mESC. We would like to thank Frédéric Langevin for assistance with the derivation of mESCs. We would like to thank Gonçalo Oliveira for assistance with the mESC culture and for technical help. We would like to thank Pranay Shah and Ross Hill for assistance with the quantification of PGCs. We would like to thank Julian E. Sale, Juan Garaycoechea and members of the Crossan laboratory for critically reading the manuscript and for useful discussions. J.R and G.P.C. are supported by the Medical Research Council. This work was supported by the Medical Research Council as part of UK Research and Innovation (file reference no. MC_UP_1201/18).

## Author contributions

Conceptualisation: JR and GC; Methodology: JR and GC; Investigation: JR; Visualisation: JR; Writing–Original Draft: JR and GC; Writing–Review & Editing: JR and GC; Supervision: GC; Project Administration: JR and GC; Funding Acquisition: GC.

## Conflict of interest

The authors declare that they have no conflict of interest.

## Expended View Figure legends

**Figure EV1 -.**
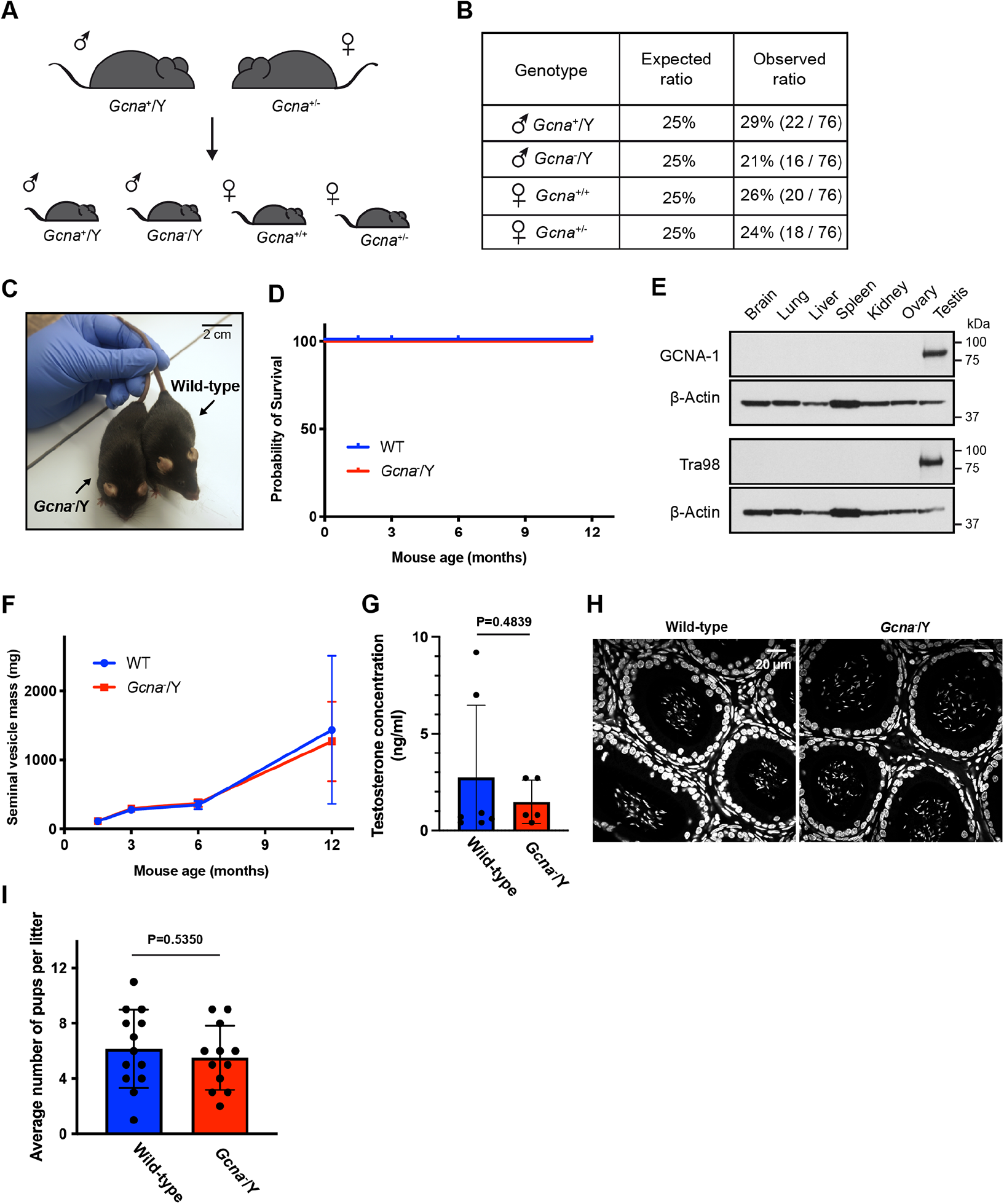
Phenotyping of GCNA-deficient male mice. A. Schematic representation of the breeding strategy to obtain willd-type or *Gcna*^−^/Y males and wild-type or heterozygous females mice. B. Expected and observed frequencies of each genotypes obtained from the breeding strategy presented in (A). C. Representative photograph of 1 year old wild-type and *Gcna*^−^/Y male mice. D. Kaplan-Meier curves of the survival of wild-type and *Gcna*^−^/Y male mice up to 1 year old. 1.5 months WT (n=6 mice) and *Gcna*^−^/Y (n=5 mice), 3 months WT (n=8 mice) and *Gcna*^−^/Y (n=6 mice), 6 months WT (n=7 mice) and *Gcna*^−^/Y (n=6 mice), 12 months WT (n=9 mice) and *Gcna*^−^/Y (n=6 mice). E. Analysis of GCNA expression in different adult mice tissues by Western blot. Twenty-five micrograms of proteins were loaded per well. Blots were probed with two antibodies directed against mouse GCNA (Tra98 and GCNA-1) and with an anti-beta-actin antibody. F. Seminal vesicle weight of wild-type and *Gcna*^−^/Y male mice between 1.5 months and 12 months. 1.5 months WT (n=6 mice) and *Gcna*^−^/Y (n=5 mice), 3 months WT (n=6 mice) and *Gcna*^−^/Y (n=4 mice), 6 months WT (n=7 mice) and *Gcna*^−^/Y (n=6 mice), 12 months WT (n=8 mice) and *Gcna*^−^/Y (n=5 mice). Data represent the mean and S.D.. G. Testosterone concentration in serum of 1 year old wild-type (n=7) and *Gcna*^−^/Y (n=5) mice. Data represent the mean and S.D.. *P* value was calculated by using an unpaired t-test. H. Representative micrographs of DAPI stained epididymis sections of 6 weeks old wild-type and *Gcna*^−^/Y male mice. I. Assessment of the fertility of wild-type and *Gcna*^−^/Y male mice. Wild-type and *Gcna*^−^/Y male mice were bred with wild-type females and the number of alive pups were counted after birth. Wild-type (n=3 mice, 13 litters) and *Gcna*^−^/Y (n=3 mice, 12 litters). Data represent the mean and S.D.. *P* value was calculated by using an unpaired t-test.

**Figure EV2 -.**
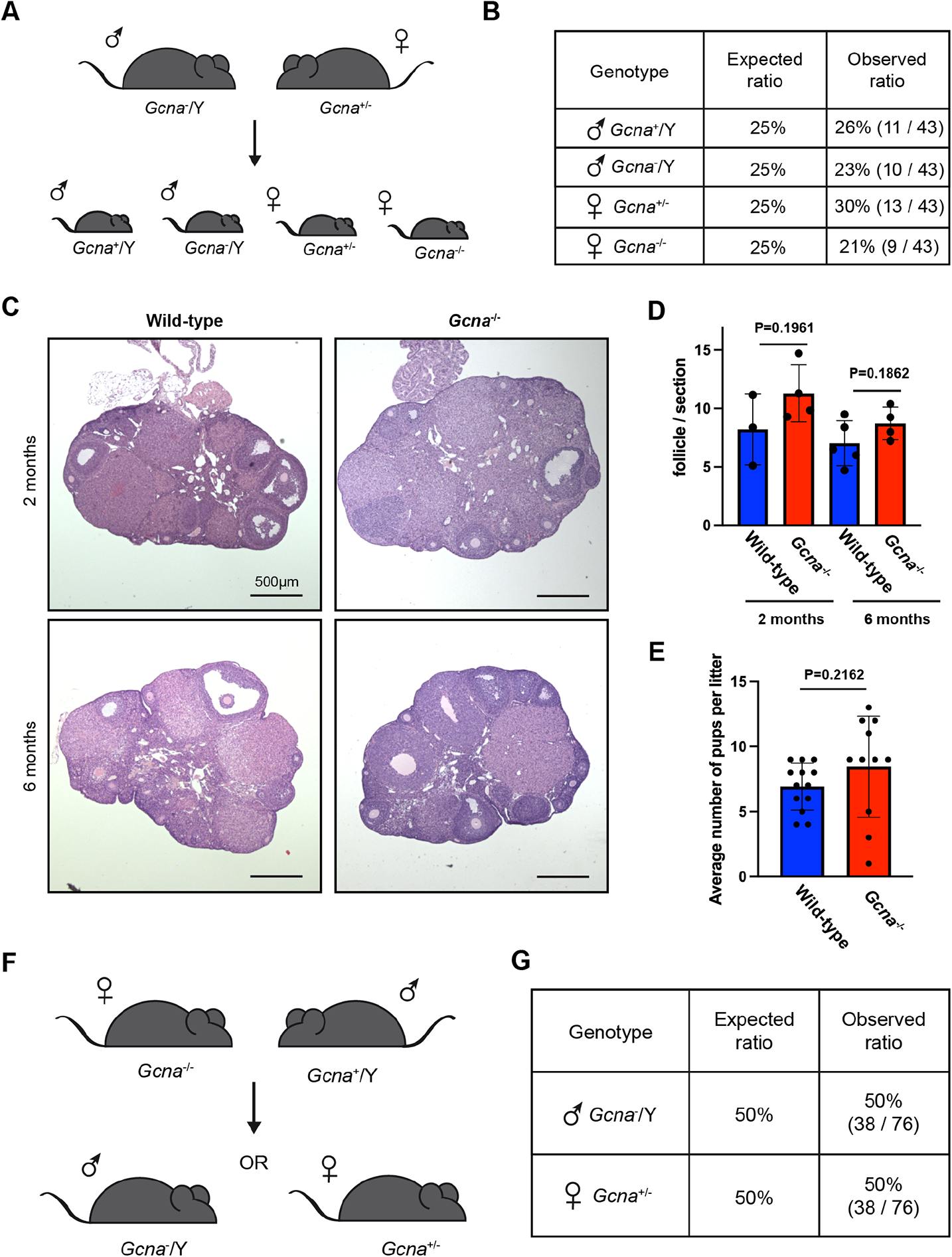
Fertility of GCNA-deficient female mice. A. Schematic representation of the breeding strategy to obtain wild-type or *Gcna*^−^/Y males and heterozygous and *Gcna*^−/-^ females mice. B. Expected and observed frequencies of each genotypes obtained from the breeding strategy presented in (A). C. Micrographs of Haematoxylin and Eosin stained ovaries sections of 2 and 6 months old wild-type and *Gcna*^−/-^ mice. D. Quantification of follicles per ovaries section of 2 and 6 months old mice. Two months WT (n=3 mice) and *Gcna*^−/-^ (n=4 mice), 6 months WT (n=5 mice) and *Gcna*^−/-^ (n=4 mice). Data represent the mean and S.D.. *P* values were calculated by using an unpaired t-test. E. Assessment of the fertility of wild-type and *Gcna*^−/-^ female mice. Wild-type and *Gcna*^−/-^ female mice were bred with wild-type males and the number of alive pups were counted after birth. Wild-type (n=3 mice, 13 litters) and *Gcna*^−/-^ (n=3 mice, 11 litters). Data represent the mean and S.D.. *P* values were calculated by using an unpaired t-test. F. *Gcna*^−/-^ female mice were bred with wild-type males and pups were genotyped after birth. G. Expected and observed frequencies of each genotypes obtained from the breeding strategy presented in (F).

**Figure EV3 -.**
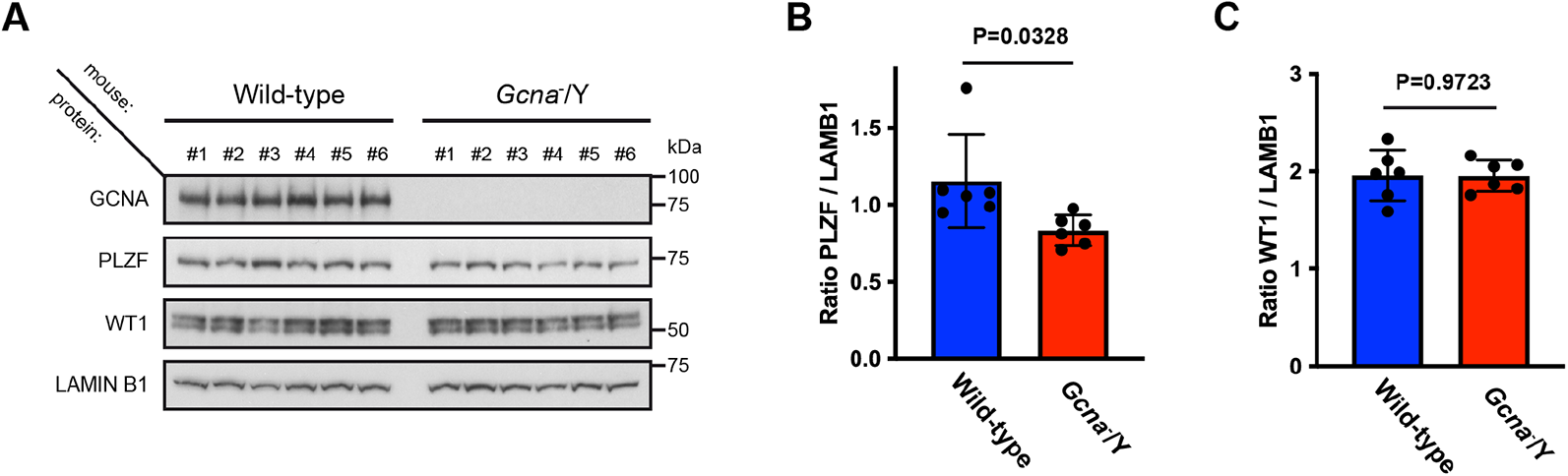
Decrease of PLZF expression in testes of 10 weeks old GCNA-deficient mice. A. Analysis of PLZF and WT1 expression in testes of 10 weeks old mice by Western blot. Twenty-five micrograms of proteins per wells were loaded. Blot was probed with antibodies directed against mouse GCNA (GCNA-1), PLZF, WT1 and with an anti-LAMIN B1 antibody. B. Quantification of the PLZF signal in (A) relative to LAMIN B1 (wild-type, n=6 mice and *Gcna*^−^/Y, n=6 mice). Data represent the mean and S.D.. *P* value was calculated by using an unpaired t-test. C. Quantification of the WT1 signal in (A) relative to LAMIN B1 (wild-type, n=6 mice and *Gcna*^−^/Y, n=6 mice). Data represent the mean and S.D.. *P* value was calculated by using an unpaired t-test.

**Figure EV4 -.**
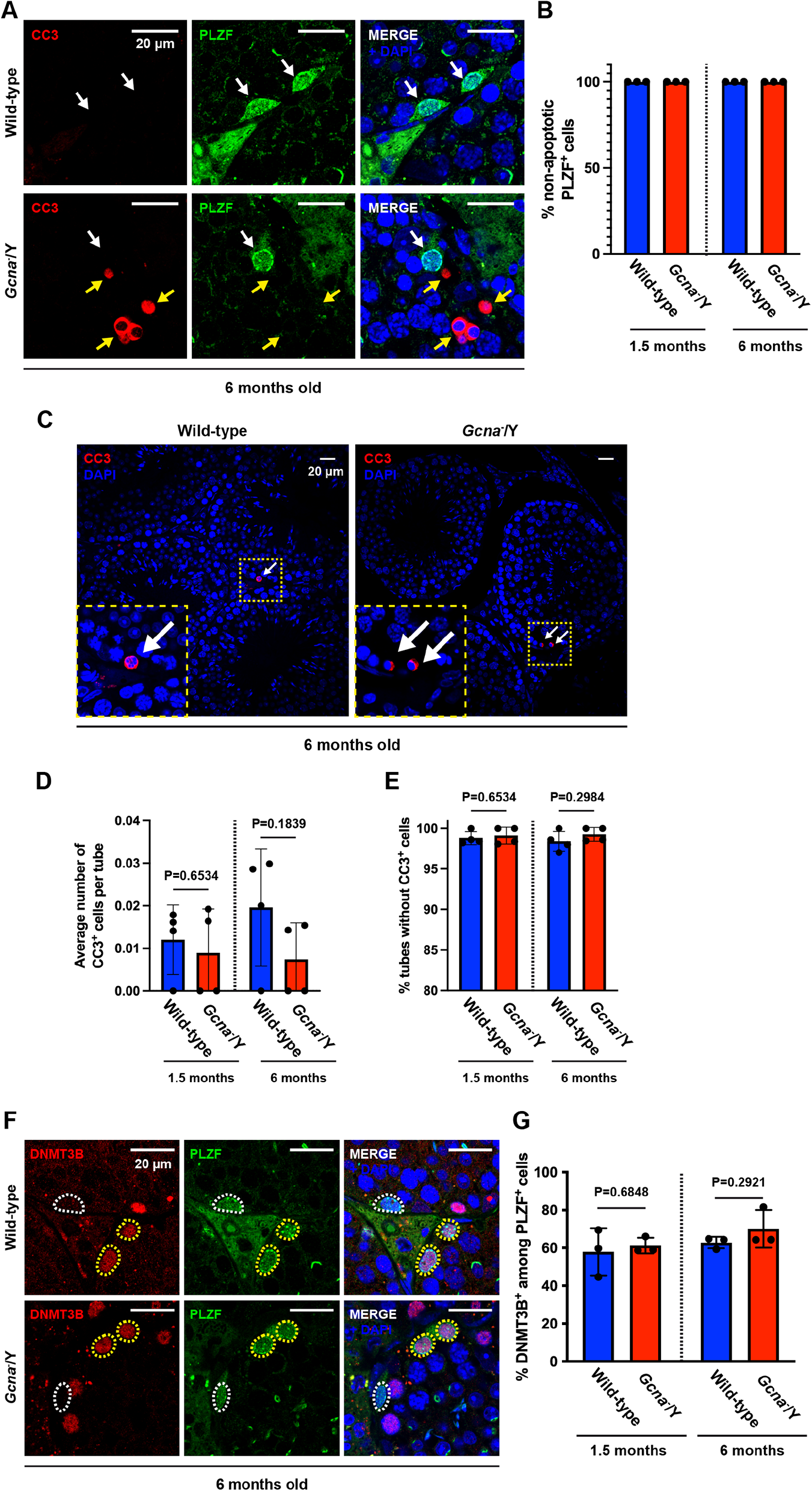
SSCs are not exhibiting increased apoptosis nor enhanced differentiation in absence of GCNA. A. Immunofluorescence staining of wild-type and *Gcna*^−^/Y testis sections from 6 months old mice. Cleaved Caspase 3 (CC3) was displayed in red, PLZF in green and DNA (DAPI stained) in blue. White arrows highlight PLZF positive SSCs and yellow arrows highlight CC3 positive germ cells. B. Frequency of PLZF positive cells exhibiting a CC3 positive signal at 1.5 and 6 months old. A minimum of 50 PLZF positive cells are scored per mouse. Data represent the mean and S.D.. N=3 mice for each genotype and age. C. Immunofluorescence staining of cleaved caspase 3 (CC3) on testis sections of 6 months old wild-type and *Gcna*^−^/Y mice. D. Average number of CC3 positive cells per seminiferous tubes at 6 months old. At least 50 seminiferous tubes are scored per mouse. Data represent the mean and S.D.. N=3 mice for each genotype. E. Frequency of seminiferous tubes that are not exhibiting any CC3 positive cells in 6 months old mice. At least 50 seminiferous tubes are scored per mouse. Data represent the mean and S.D.. N=3 mice for each genotype. F. Immunofluorescence staining of wild-type and *Gcna*^−^/Y testis sections from 6 months old mice. DNMT3B was displayed in red, PLZF in green and DNA (DAPI stained) in blue. Cells positive for both PLZF and DNMT3B are highlighted in yellow and PLZF only cells are highlighted in white. G. Frequency of PLZF positive cells also positive for DNMT3B at 1.5 and 6 months old. A minimum of 50 PLZF positive cells are scored per mouse. Data represent the mean and S.D.. N=3 mice for each genotype and age.

**Figure EV5 -.**
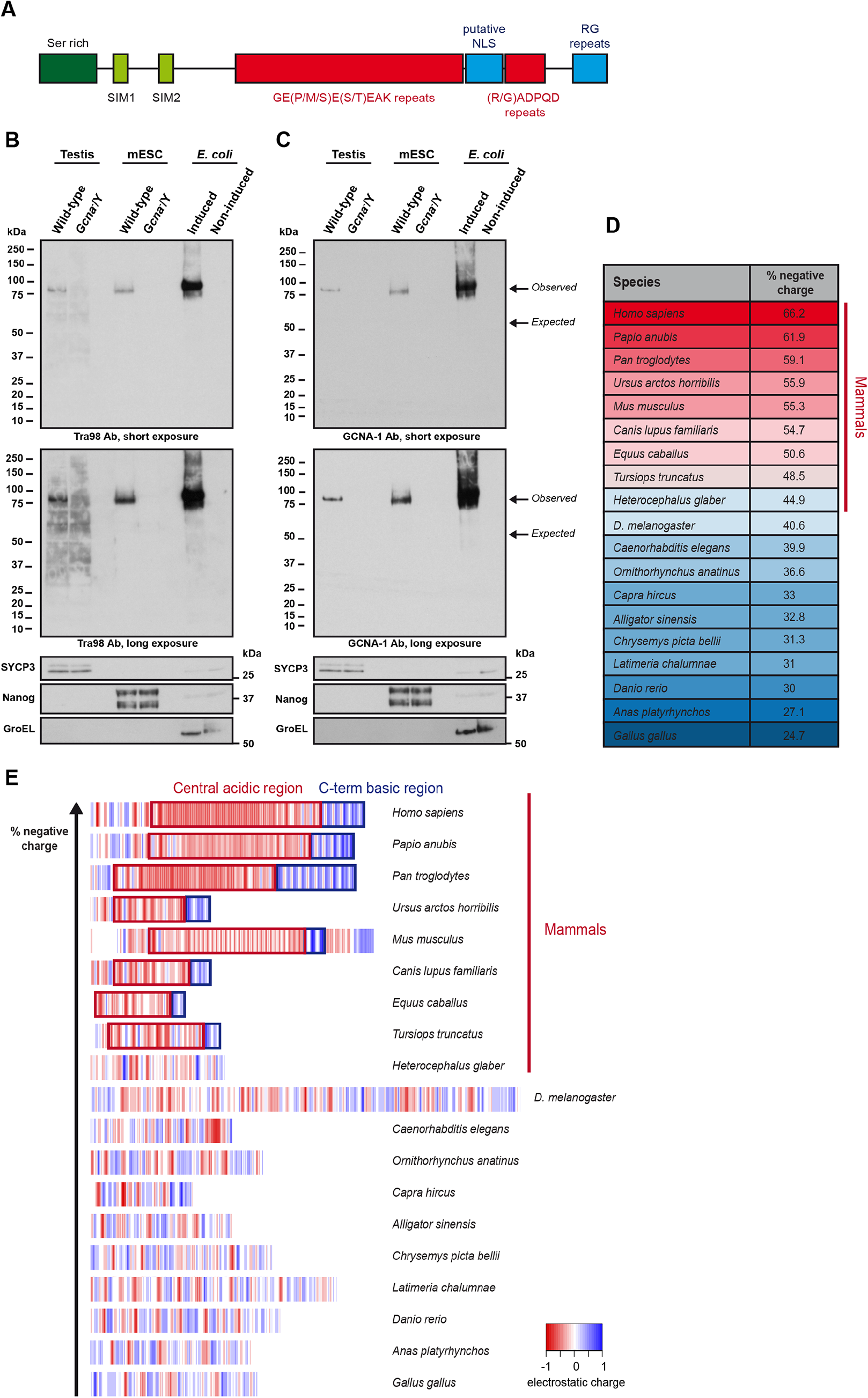
Mouse GCNA is related to the Intrasically Disordered Region (IDR) of human GCNA. A. Schematic representation of the annotated mouse GCNA protein. Mouse GCNA is fully disordered and possess multiple repeats domains. Among them, two are highly acidic (red) and two are basic (blue). SUMO-interacting motifs are represented in light green. A serine-rich region is represented in dark green. B. Western blot of GCNA from different sources. This Western blot displays extracts from wild-type and *Gcna*^−^/Y testes, wild-type and *Gcna*^−^/Y mESCs and *E.coli* transformed with an expression plasmid (induction with IPTG). Then, the blot was probed against mouse GCNA (TRA98), a testis marker (SYCP3), a mESC marker (Nanog) and a marker for *E.coli* (GroEL). Data is representative from two independent experiments. C. Western blot of GCNA from different sources. This Western blot displays extracts from wild-type and *Gcna*^−^/Y testes, wild-type and *Gcna*^−^/Y mESCs and *E.coli* transformed with an expression plasmid (induction with IPTG). Then, the blot was probed against mouse GCNA (GCNA-1), a testis marker (SYCP3), a mESC marker (Nanog) and a marker for *E.coli* (GroEL). Data is representative from two independent experiments. D. Percentage of negative charges in IDRs of metazoans GCNA proteins. Proteins are distributed according to their percentage of negative charges. E. Distributions of charges in IDRs of metazoan GCNA proteins. Acidic residues are displayed in red and basic residues in blue. Proteins are distributed according to their percentage of negative charges.

## Notes

### Competing Interest Statement

The authors have declared no competing interest.

