## Appendix PDF for "GCNA is a histone binding protein required for spermatogonial stem cell maintenance"

### TABLE OF CONTENT

This appendix contains 8 figures.

### APPENDIX FIGURES


**Appendix Figure S1** page 3

**Appendix Figure S2** page 4

**Appendix Figure S3** page 5

**Appendix Figure S4** page 6

**Appendix Figure S5** page 7

**Appendix Figure S6** page 8

**Appendix Figure S7** page 9

**Appendix Figure S8** page 10

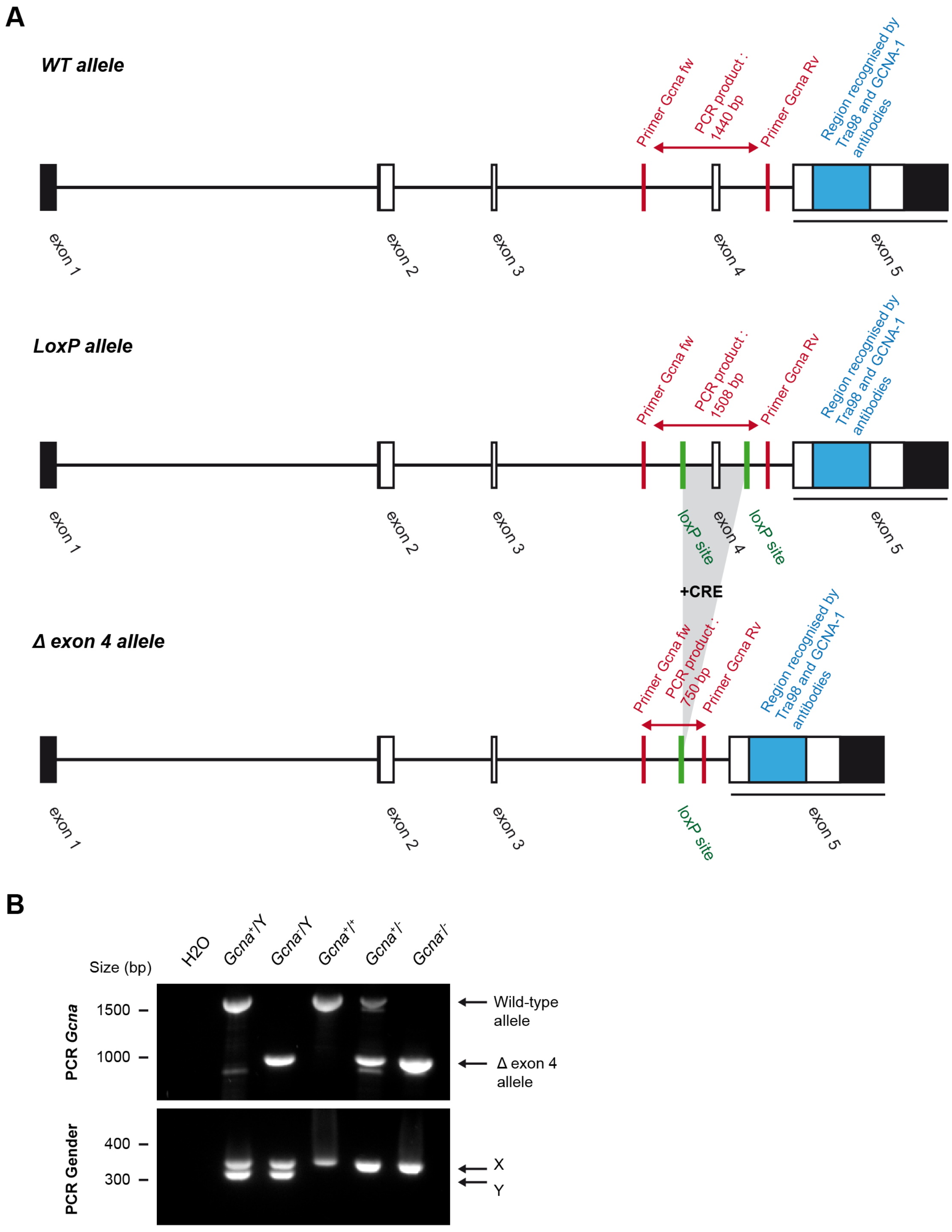


**Appendix Figure S1 - Generation and validation of GCNA-deficient mice.**

A. Schematic representation of the different alleles of Gcna used for this study. Coding regions in exons are displayed in white boxes and non-coding are displayed in black boxes. Primers and PCR fragments lengths used for genotyping are highlighted in red. LoxP sites are highlighted in green. The region recognised by two antibodies directed against mouse GCNA (Tra98, GCNA-1) is highlighted in blue.

B. Endpoint genotyping PCRs obtained using the PCR primers showed in (A).

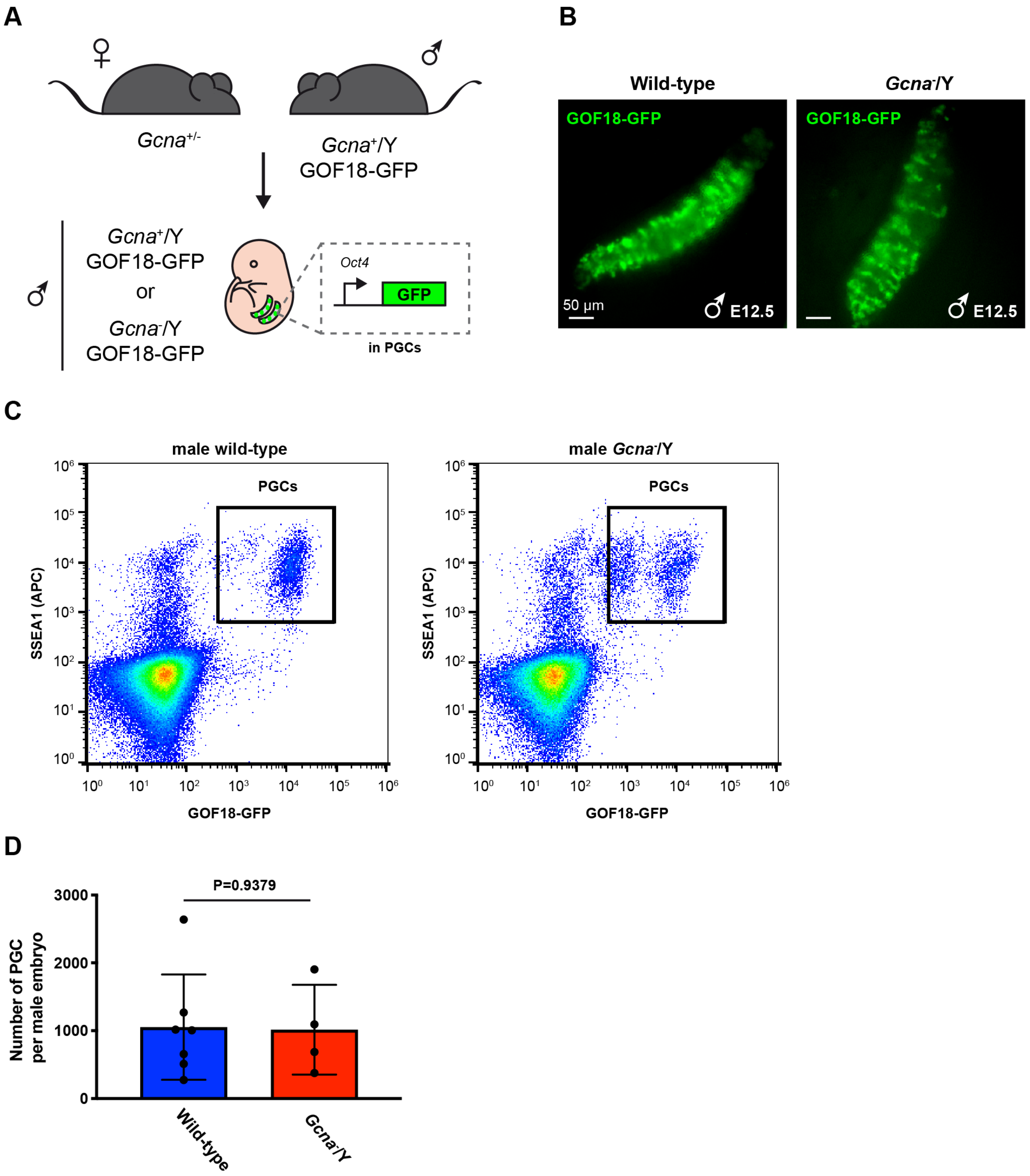


**Appendix Figure S2 - GCNA is dispensable for male PGC development.**

A. Schematic representation of the breeding strategy to obtain wild-type or *Gcna*-/Y male carrying the GOF18-GFP PGC reporter.

B. Micrographs of GOF18-GFP fluorescence in E12.5 male gonads from wild-type and *Gcna*-/Y embryos carrying the GOF18-GFP PGC reporter.

C. Representative flow cytometry plots (SSEA1/GOF18-GFP) from E12.5 wild-type and *Gcna*-/Y male embryos.

D. Quantification of PGCs (SSEA1 and GOF18-GFP double positive cells) from male embryos at E12.5 (wild-type, n=7 embryos and *Gcna*-/Y, n=4 embryos). Data represent the mean and S.D.. P value was calculated by using an unpaired t-test.

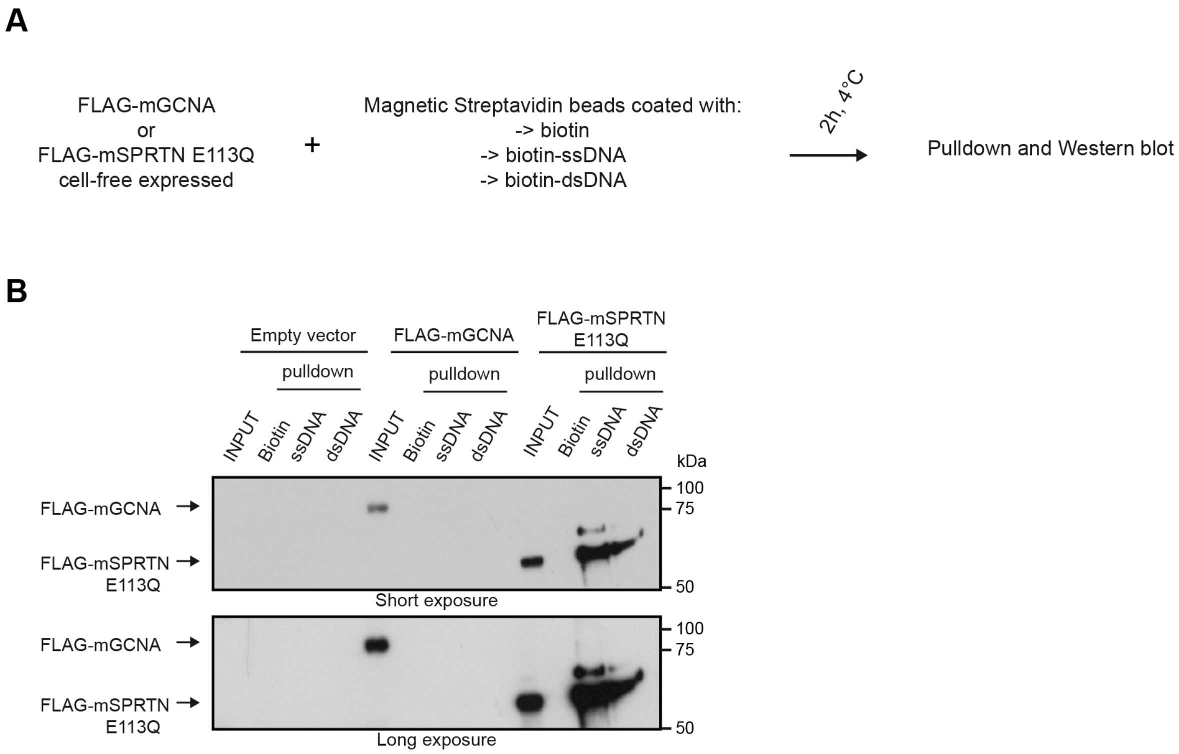


**Appendix Figure S3 - Mouse GCNA does not bind DNA in vitro.**

A. A DNA pulldown was performed with recombinant FLAG-mGCNA and FLAG-mSPRTN E113Q expressed in a cell free system. Products of this pulldown were then analysed by Western blot.

B. Western blot of the DNA pulldown. The blot was probed with an anti-FLAG antibody. Data is representative from two independent experiments.

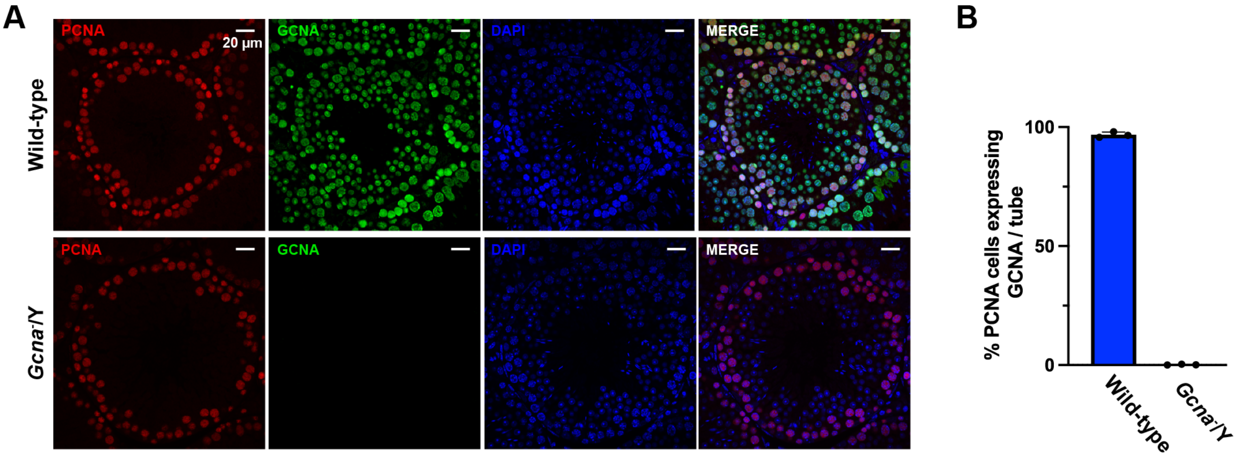


**Appendix Figure S4 - Co-expression of PCNA and GCNA in mouse pre-spermatid cells.**

A. Immunofluorescence staining of wild-type and *Gcna*$^{-}$/Y adult testis sections. PCNA was displayed in red, GCNA (Tra98) in green and DNA (DAPI stained) in blue. Data is representative from three independent experiments.

B. Frequency of PCNA positive cells co-expressing GCNA per tube. A minimum of 580 cells are scored per mouse. N=3 mice for each genotype. Data represent the mean and S.D.. P value was calculated by using an unpaired t-test.

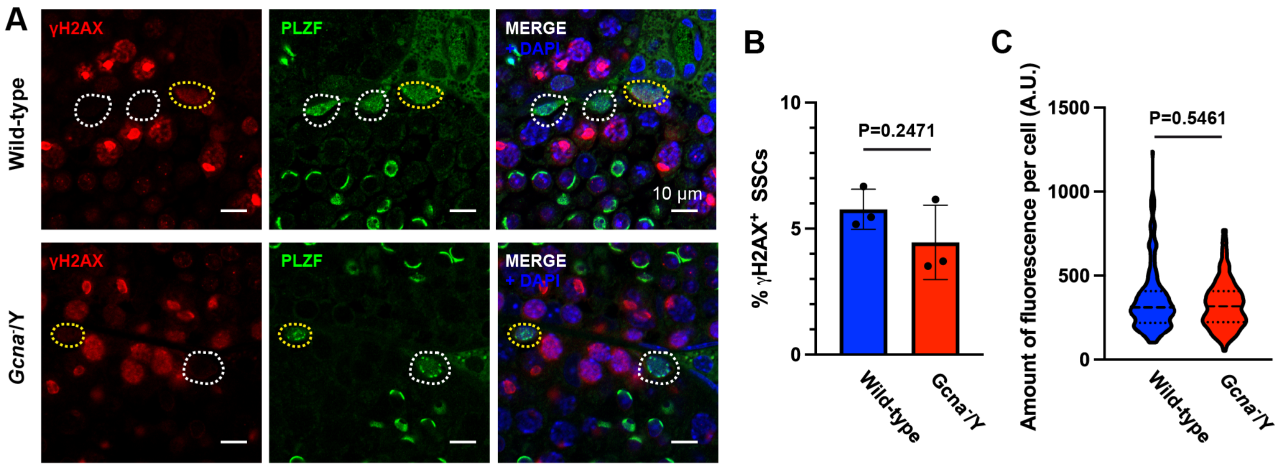


**Appendix Figure S5 - DNA breaks in SSCs of 6 weeks old mice.**

A. Immunofluorescence staining of wild-type and *Gcna*$^{-}$/Y testis sections from 6 weeks old mice. γH2AX was displayed in red, PLZF in green and DNA (DAPI stained) in blue. Cells positive for both PLZF and γH2AX are highlighted in yellow and PLZF only cells are highlighted in white.

B. Frequency of PLZF positive cells also positive for γH2AX at 6 weeks old. A minimum of 50 PLZF positive cells are scored per mouse. Data represent the mean and S.D.. N=3 mice for each genotype. P values were calculated by using an unpaired t-test.

C. Quantification of the amount of fluorescence (integrated density) of γH2AX per SSC, in wild-type and *Gcna*$^{-}$/Y testis from 6 weeks old mice. Wild-type (N=179 SSCs from three mice), *Gcna*$^{-}$/Y (N=178 SSCs from three mice). Data represent the median and interquartile range. P values were calculated by using a two-tailed Mann-Whitney test.

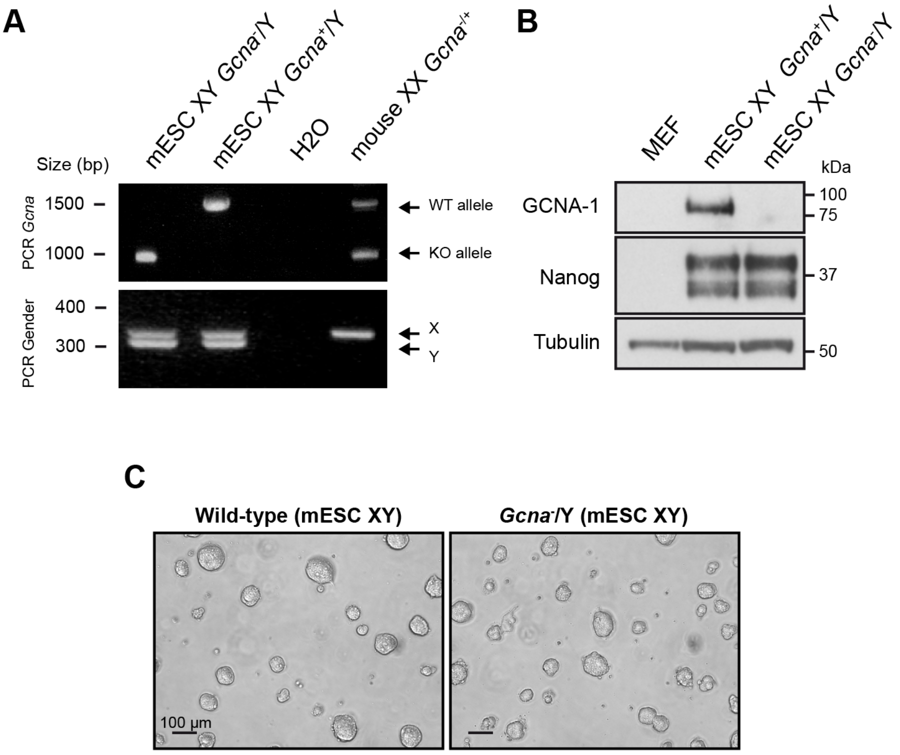


**Appendix Figure S6 - Generation and validation of GCNA-deficient mESC.**

A. Validation of the mouse ES cell lines by PCR, by using the PCR primers showed in Appendix Figure S1A.

B. Validation of the mouse ES cell lines by Western blot. Blot was probed with an antibody directed against mouse GCNA (GCNA-1), with an anti-Nanog antibody and an anti-tubulin antibody.

C. Micrographs of the mouse ES cell lines in culture.

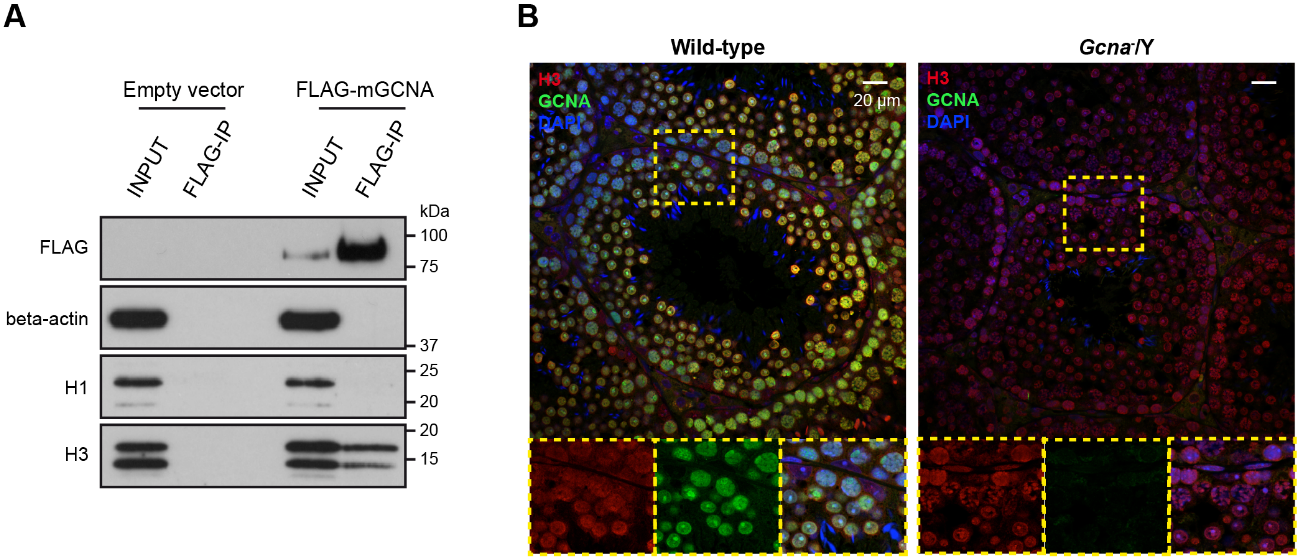


**Appendix Figure S7 - Biochemical features of mouse GCNA.**

A. Western blot of the FLAG immunoprecipitation. 3T3 cells were transiently transfected with FLAG-mGCNA and a FLAG immunoprecipitation was performed on the soluble fraction. The blot was probed with anti-FLAG, anti-H1, anti-H3 and anti-beta-actin antibodies. Data is representative from two independent experiments.

B. Immunofluorescence staining of wild-type and *Gcna*$^{-}$/Y adult testis sections. H3 was displayed in red, GCNA in green and DNA (DAPI stained) in blue. Bottom panels are magnifications of regions highlighted by the yellow boxes.

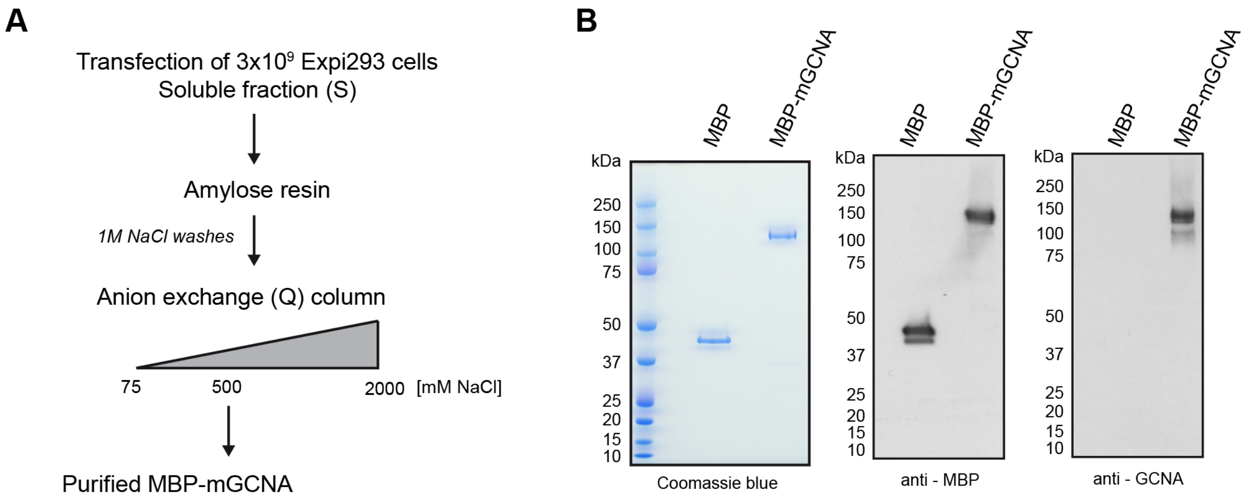


**Appendix Figure S8 - Purification of recombinant MBP-mGCNA.**

A. Scheme of purification strategy.

B. Left panel: Coomassie staining of purified MBP-mGCNA and MBP analysed by SDS-PAGE. Middle and right panels: Western blot analyses of purified MBP-mGCNA and MBP, respectively stained with anti-MBP and anti-GCNA (GCNA-1) antibodies.
